# Effects of rescaling forward-in-time population genetic simulations

**DOI:** 10.1101/2025.04.24.650500

**Authors:** Jacob I. Marsh, Sachin Kaushik, Parul Johri

## Abstract

Forward-in-time population genetic simulations enable modelling of a wide array of complex evolutionary scenarios. Simulating small genomic regions, and rescaling by reducing population size by a scaling factor while maintaining population-scaled parameters, are common approaches for improving computational tractability. However, simulating whole chromosomes in large populations remains computationally prohibitive and it’s unclear what scaling factors produce consistent evolutionary dynamics between rescaled and unscaled populations. Here, we thoroughly test the effects of rescaling in populations experiencing direct and linked effects of selection for various scaling factors and genomic lengths in *D. melanogaster*-like populations by comparing them with theoretical expectations. We find that while rescaling has minimal effects with most length and scaling factor combinations, simulating long regions with large scaling factors can introduce biases in summary statistics including neutral diversity, allele frequencies, linkage disequilibrium, and non-neutral divergence. These deviations occur particularly when the crossover rate exceeds 0.44 per individual/generation because substantial multiple-crossover events are likely to occur within individuals, reducing the effective rate of genetic recombination. In addition, when the genome-wide deleterious mutation rate is high (*U* ≫ 1) in highly rescaled long regions, we observe increased Hill-Robertson interference effects and progeny skew, the extent of which was strongly dependent on the fitness effects of selected mutations. We find that hitchhiking effects near functional regions are relatively unaffected by rescaling the population, even with large scaling factors. Our findings expose potential pitfalls when simulating long regions with rescaling and highlight parameter spaces within which expected evolutionary dynamics are conserved.

## Introduction

Forward-in-time simulations are increasingly important tools for modelling complex population genetic dynamics. Forward-in-time simulators as opposed to coalescent (backward-in-time) simulators (*e.g.,* Excoffier et al. 2021; Baumdicker et al. 2022) can more effectively capture complex population genetic dynamics such as selection. Forward-in-time simulation software such as *SLiM* (Haller and Messer 2023; see also Guillaume and Rougemont 2006; Hernandez 2008; Thornton 2019; Matthey-Doret 2021) and *fwdpp/fwdpy* (Thornton 2014; Thornton 2019) provides a flexible framework for modelling complex populations with specific genome architectures. This allows researchers to set parameters (controlling *e.g.*, recombination, mutations, selection, population history) informed by real populations or systems of interest to generate expected sequence variation data.

Forward-in-time simulations have been used in research to, for example, test the effects of selection on population genomic inference (*e.g.,* Nam et al. 2017; Pouyet et al. 2018; Booker 2020; Huang et al. 2021; Johri, Riall, et al. 2021; Daigle and Johri 2024; Marsh and Johri 2024). However, given that many real populations have large effective population sizes and large genomes, researchers typically need to simulate small chromosomal regions and/or rescale population parameters to achieve computational tractability. Thus, to accurately model and measure the linked effects of selection acting across the genome, simulating longer chromosomal regions is required. In addition, simulation-based inference approaches use simulated data as input to infer population parameters with statistical approaches such as approximate Bayesian computation (ABC; Beaumont et al. 2002) and by training machine learning models (Schrider and Kern 2018; Korfmann et al. 2023). Simulation-based inference methods require testing of a wide parameter space and accuracy can be limited by bottlenecks in the computational load involved with the large number of simulations. For such applications, there is a pressing need to better understand the limits of rescaling when simulating regions of different chromosomal lengths.

Rescaling forward-in-time simulations in a panmictic population involves reducing the effective population size of the unscaled population (*N*_*e*_) by a scaling factor (*X*), while increasing the mutation rate (μ), recombination rate (*r*), and selective strength (*s*) by the same factor (see Methods), as demonstrated first by Robertson (1960), as well as in Hill and Robertson (1966). Population dynamics are typically expected to be stable as long as the population-scaled parameters are consistent with the unscaled population (*e.g.,* N_scaled_ × μ_scaled_ = N_e_ × μ; McVean and Charlesworth 2000; Hoggart et al. 2007; Kim and Wiehe 2008; Kaiser and Charlesworth 2009), and parameters do not violate diffusion limits (Norman 1975; Wakeley 2005; Barton and Otto 2005; Innan and Sakamoto 2021). However, Uricchio and Hernandez (2014) found that rescaling could skew patterns of diversity in populations experiencing recurrent selective sweeps by increasing the rate of interference between selected alleles. In addition, recent findings by Dabi and Schrider (2024) suggest that rescaling can significantly bias fixation times, fixation probabilities, allele frequencies, and linkage disequilibrium in their simulated scenarios. Ferrari *et al*. (2024) recently showed that increasing the length of the simulated region (*L*) in rescaled simulations can lead to reduced nucleotide diversity and a shift toward fewer intermediate-frequency alleles. Although these studies have demonstrated several important risks of rescaling, three issues remain unclear: (a) When is rescaling reasonably accurate and when might it no longer effectively model the unscaled population? (b) Why does rescaling in certain scenarios lead to the observed biases? Finally (c), can we theoretically predict the effect of rescaling on some population genetic quantities?

As chromosome-wide crossover rates approach and exceed 1 due to rescaling (*e.g.*, Haldane 1919), there is a non-linear increase in the effective rate of recombination with the scaling factor, likely resulting in the observed biases. Here, we test different combinations of the scaling factor and the length of the simulated region (which will determine how quickly we approach this limit) to see how rescaling impacts the direct and linked effects of purifying and positive selection. We examine diversity (π) at neutral sites, divergence, the site frequency spectrum (SFS), and linkage disequilibrium (LD) for simulated regions of different scaling factors and lengths. Following that, we investigate the impacts of rescaling on local effects of selection at linked neutral sites near functional regions experiencing recurrent selection, and post-fixation of beneficial mutations. Moreover, we compare the linked effects of selection on neutral diversity to that expected theoretically with rescaling, both under assumptions of a linear recombination map, and accounting for the increased incidence of multiple crossovers due to rescaling to ask if such effects are theoretically predictable. We do so in populations simulated with parameters similar to *D. melanogaster* populations because not only has *D. melanogaster* been an important model organism in population genetics, but their large populations require drastic rescaling to simulate in a reasonable amount of time. Finally, we provide recommendations for effective rescaling across different simulation parameters and discuss the processes that cause biases due to excessive rescaling.

## Methods

### Simulations of genomic regions

Simulations were performed using *SLiM* 4.0.1 (Haller and Messer 2023). Wright-Fisher *Drosophila melanogaster*-like populations were simulated under a constant effective population size, *N*_*e*_ = 1 × 10^6^ diploid individuals, a mean mutation rate of 3 × 10^−9^ per site/generation (Keightley et al. 2014), and a mean sex-averaged crossover rate of 1 × 10^−8^ per site/generation (Comeron et al. 2012), prior to rescaling. We varied the length (*L*) of the simulated region from 10 kb to 25 Mb and tested a range of scaling factors (*X*) from 25 to 625, where scaled population sizes were reduced below 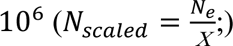, and scaled mutation and recombination rates were increased by the same factor (*μ_scaled_* = *X* × 3 × 10^−9^; *r*_*scaled*_ = *X* × 1 × 10^−8^). Time was decreased by the same factor so that the burn-in was implemented for 14*N*_*scaled*_ generations (instead of 14*N*_*e*_ generations in an unscaled simulation). The selection coefficient was also increased by the same factor so that the population scaled strength of selection remains constant (2*Ns* = 2*N*_*scaled*_ *s*_*scaled*_). Genome architecture was not modelled in these simulations, *i.e.,* mutation and recombination rates were constant across sites and selected mutations were uniformly distributed across the simulated region. Ten replicates were simulated for each parameter combination of length and scaling factor. Results from parameter combinations for which all ten replicates did not finish during the 14*N*_*scaled*_ generation burn-in were not included (see *SLiM* scripts at: https://github.com/JohriLab/Simulation_Rescaling/tree/main/slim).

#### Simulating fitness effects

Half of all new mutations in the simulated region experienced selection and all mutations were assumed to be semidominant (ℎ = 0.5) unless otherwise specified. Note that we use “*s*” to represent the selective advantage of the mutant homozygote relative to wildtype, i.e., positive values indicate beneficial alleles and negative values correspond to deleterious ones. In order to test the effects of background selection (BGS), purifying selection was simulated with varying parameters: strong purifying selection with constant selective effects (2*Ns* = −100), weak purifying selection with constant selective effects (2*Ns* = −20), purifying selection generated by dominant lethal mutations with constant selective effects (*s* = −1; ℎ = 1), and purifying selection with selective effects drawn from a distribution of fitness effects (DFE) estimated by Johri et al. (2020). The DFE was comprised of three non-overlapping uniform distributions representing the weakly deleterious (−1 ≥ *Ns* > −10), moderately deleterious (−10 ≥ 2*Ns* > −100), and strongly deleterious (−100 ≥ 2*Ns* > −2*N*_*scaled*_) classes of mutations, comprising 65.33%, 5.33%, and 29.33% of all selected mutations respectively.

#### Simulating recurrent selective sweeps

When simulating recurrent selective sweeps due to beneficial mutations, an *f*_*pos*_ proportion (0.002) of new selected mutations was assumed to be beneficial (as estimated by Campos *et al.,* 2017), while 1 − *f*_*pos*_ were modelled as strongly deleterious with constant selective effects (2*Ns* = −100). Beneficial mutations were similarly assumed to be semidominant (ℎ = 0.5), and followed an exponential DFE with 2*Ns*_*adv*_ = 250, where 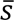 is the mean, reflecting parameters estimated by Campos *et al*. (2017).

#### Gene conversion

Gene conversion was not modelled, except in specific simulations, in order to better isolate and understand the impacts of other factors (notably selection and recombination) when rescaling is present. When testing the effects of gene conversion on diversity, gene conversion was modelled in *SLiM* using parameters estimated by Miller *et al*. (2016): the sex-averaged gene conversion initiation rate was set to *g*_*c*_ = *X* × 1 × 10^−8^, while the recombination rate was adjusted so that the crossover rate remained unchanged; mean tract length was set to 440 bp (invariant to scaling factor), with only simple conversions modelled.

#### Simulations with constant *U* or *R*

While simulated crossover and mutation rates were generally modelled as linearly increasing with scaling factor, in order to test biases due to rescaling with alternative ratios of *U* and *R* as may be relevant to other species, we tested scenarios with varying rates of region-wide crossover rate (*R* = *r*_*scaled*_ × *L*) and deleterious mutation rate per diploid individual (*U* = 2μ_*scaled*_ × *L*). For these simulations, the per-site crossover or mutation rate was adjusted so that the mean region-wide values remained fixed: constant *R* = 0.5, or constant *U* = 0.5, regardless of scaling factor. In all other respects, these simulations were modelled identically to the simulations of genomic regions experiencing strong purifying selection with constant selective effects as described above.

### Simulations evaluating the recovery of neutral diversity away from a selected region

A selected region of fixed length (10 kb) was simulated with a 30 kb flanking neutral region (see Figure 5). The selected region experienced only purifying selection (DFE from Johri *et al.,* 2020), or both purifying and recurrent positive selection (*f*_*pos*_ = 0.002; 2*Ns*_*adv*_ = 250). These simulations were performed for different scaling factors, with 10,000 replicates each. Nucleotide site diversity (π), Tajima’s *D* (Tajima 1989), and Fay and Wu’s *H* (Fay and Wu 2000) were calculated from the neutral region using derived allele counts, in 250 bp non-overlapping bins, after 14*N*_*scaled*_ burn-in using custom *R* scripts (https://github.com/JohriLab/Simulation_Rescaling/blob/main/scripts/calculate_fay_wu_h.py). Results reflect statistics averaged across the 10,000 replicates.

### Simulations evaluating the recovery of neutral diversity due to the fixation of a beneficial mutation

The recovery of neutral diversity during and after the fixation of a favorable mutation was quantified using varying scaling factors. A single selected site was simulated with a flanking neutral region of 40 kb. A new beneficial mutation (2*Ns* = 100, 2*Ns*_*adv*_ = 1000) was periodically introduced at the selected site after 14*N*_*scaled*_ generations of burn-in. If the new beneficial mutation was lost, another beneficial mutation would be introduced in the following generation. Following the fixation of a beneficial mutation, a new beneficial mutation was not introduced for the subsequent 4*N*_*scaled*_ generations. Summary statistics were calculated using *SLiM* while a new beneficial mutation increased in allele frequency, reached fixation, and at specified generations post-fixation. The results reflect statistics averaged across a minimum of 1,792 replicates; more replicates were used for plots with higher scaling factors as they required less computational resources. Figures were generated in *R* (R Core Team 2024; https://github.com/JohriLab/Simulation_Rescaling/tree/main/scripts).

### Calculating summary statistics

Mean nucleotide diversity at neutral sites (π_*neu*_) was calculated using the calcHeterozygosity function within *SLiM*, applied to only neutral mutations and multiplied by the total proportion of neutral sites in the genome. The relative change in nucleotide diversity across neutral sites due to the effects of background selection under different selection parameters was estimated by 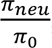, where π represents the theoretical expectation of nucleotide diversity under strict neutrality (π_0_ = 4*N*_*e*_ μ = 0.012). Local BGS and sweep effects are known to be weaker approaching the ends of a chromosome due to the reduced density of proximal selected sites (Nordborg et al. 1996; Charlesworth 2013). Therefore, we tested measures of diversity including only the central 50% of sites in the simulated region (Table S1) as well as excluding 20 kb from each end of the simulated region (Table S2). Both changes to the measured regions had negligible effects on observed diversity across the different *L* and *X* combinations (compare Table 1 to Table S1 and Table S2). Therefore, unless specified, diversity was calculated using all sites across the simulated region.

**Table 1.**
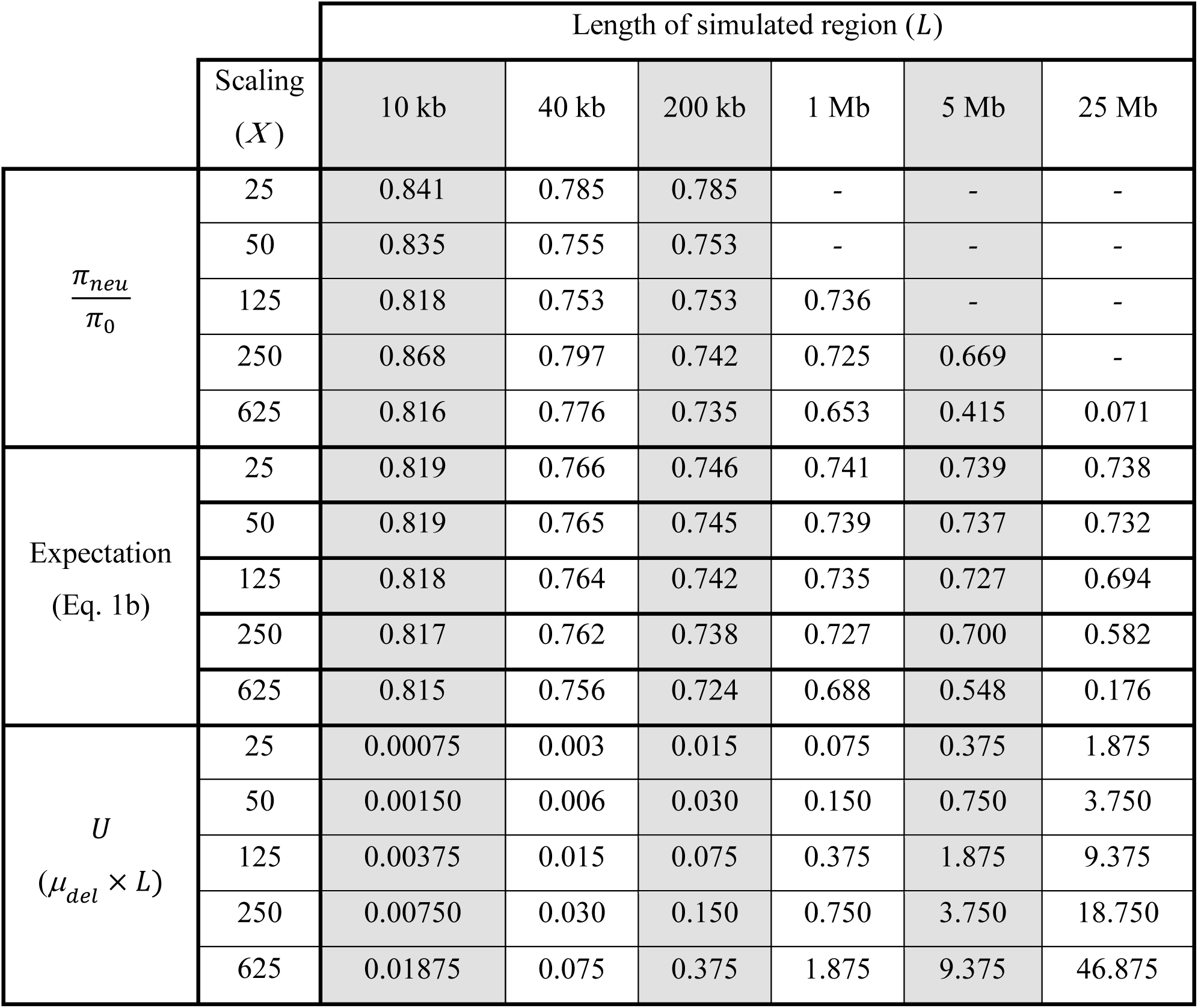
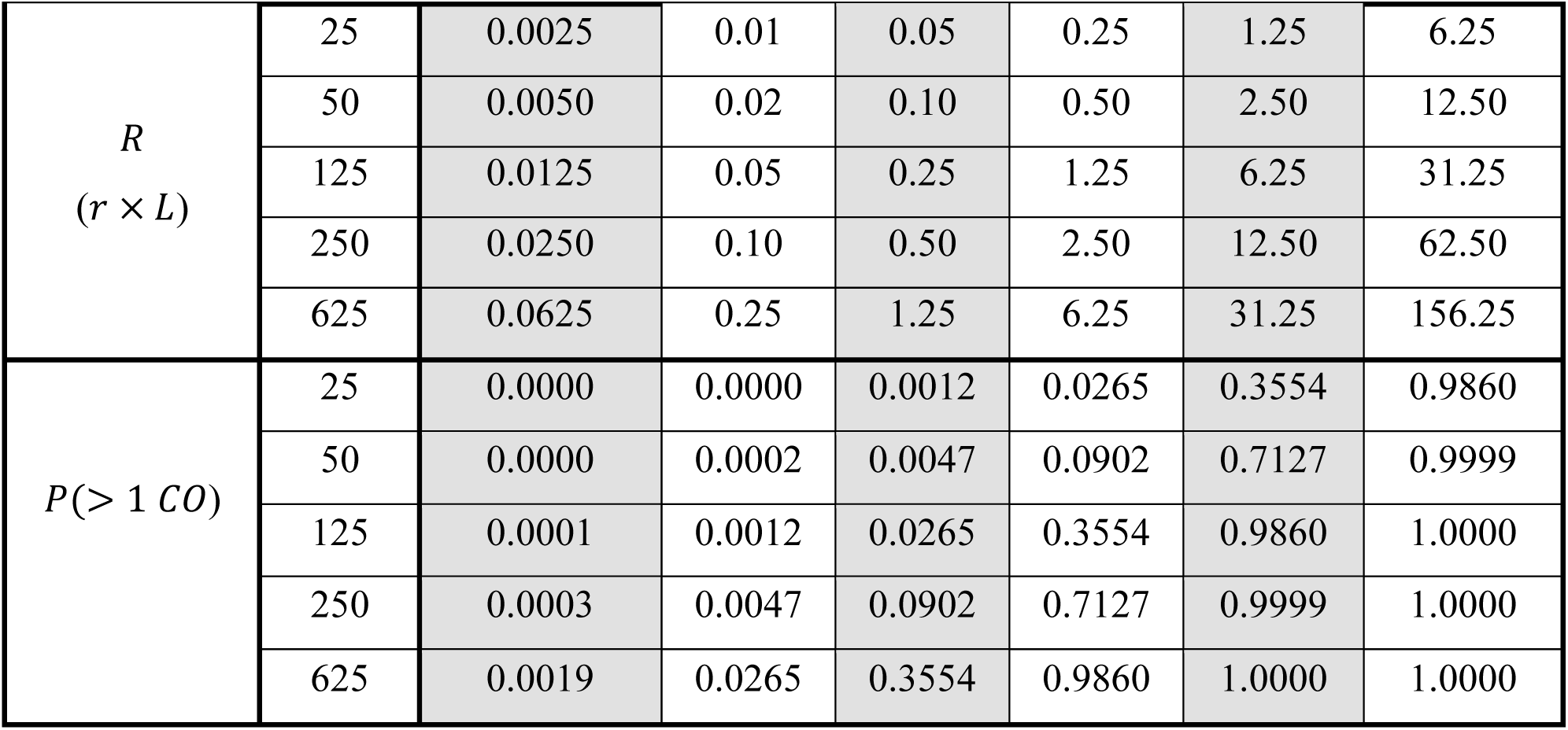
Observed nucleotide diversity at neutral sites (π_*neu*_) in simulations experiencing background selection relative to expected diversity under strict neutrality (π_0_ = 0.012), for simulations of varying length (*L*) and scaling factor (*X*). Strong purifying selection was simulated with constant population-scaled selective effects (2*Ns* = −100) applied to half of all new mutations. Corresponding simulated values of the genome-wide deleterious mutation rate, *U*, and genome-wide recombination rate per individual per generation, *R*, are shown, as well as the expected relative diversity at neutral sites due to background selection compared to under neutrality (Eq. 1b), and the probability of multiple crossover events occurring in each individual per generation, *P*(> 1 *CO*). The result from Eq. 3 was 0.7408 for all parameter combinations.

Divergence at selected sites was calculated as the proportion of all substitutions that occurred between 10*N*_*scaled*_ generations and the end of the 14*N*_*scaled*_ generations; the mean of results across replicates is displayed. Linkage disequilibrium (LD) decay was calculated between all pairwise sites with *PopLDdecay* (Zhang et al. 2019) using the *SLiM* VCF output with the following settings:-OutType 1-MAF 0-Het 1. Note that LD between a pair of SNPs separated by distance *d* is expected to remain the same between an unscaled and a rescaled population, as the expected number of crossovers between them per generation (*N*_*e*_ *rd* = *N*_*scaled*_ *r*_*scaled*_*d*) stays constant. The site frequency spectrum (SFS) for derived neutral mutations was calculated excluding invariant sites. In some simulations we tracked offspring frequencies per parental individual for each generation in the final 100 generations using ‘p1.lifetimeReproductiveOutput’ in *SLiM*. In addition, we tracked the mean and variance of fitness per individual using ‘p1.cachedFitness(NULL)’.

### Obtaining theoretical expectations of diversity due to BGS and sweeps

We calculated the nucleotide diversity at neutral sites (π_*neu*_) with BGS and selective sweeps relative to what may be expected under strict neutrality (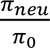, or *B* when only considering the effect of BGS) where the expectation under neutrality is provided by π_0_ = 4*N*_*e*_μ.

The following equation (Eq) predicts for a given site (*q*) the relative decrease in diversity caused by background selection (*b*_*q*_) from a single site under purifying selection *i*, where *μ_del_* is the probability that site *i* will mutate from neutral (ancestral) to a deleterious allele per generation with selective strength *s* (note that *s* will be negative for deleterious mutations), and dominance coefficient (ℎ), assuming that genetic distance can be modelled linearly by *rl*_*qi*_ (*i.e.*, no multiple crossovers) where the rate of crossovers per site per generation is *r* with physical distance *l*_*qi*_ between the focal site *q* and conserved site *i* (Nordborg et al. 1996):

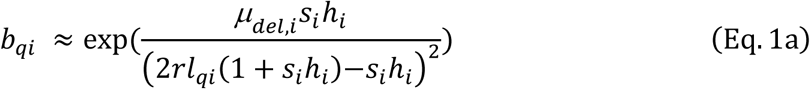

where *μ_del_*, *s* and *r* may all be linearly rescaled by a given scaling factor (*X*). However, when multiple crossovers are common and thus the frequency of recombination between *q* and *i* is not simply *rl*_*qi*_, Eq. 1a can be modified using Haldane’s mapping function (Haldane 1919) which accounts for multiple crossovers with no crossover interference effects, as is simulated in *SLiM*:

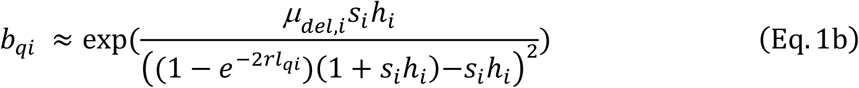

We assumed multiplicative fitness effects when considering the combined effect of all conserved sites on a given site (*B*), which we calculated as a product for all sites, so that for example *B* at the first and last position in the chromosome would be 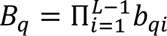. Please see https://github.com/JohriLab/Simulation_Rescaling/blob/main/scripts/ExactBcalc.R for the *R* script that implements Eq. 1a and Eq. 1b to calculate *B* given population genetic parameters as expected for our simulations. When calculating the expected *B* in simulations experiencing a discretised DFE, we sampled the DFE for each position *i* with probabilities as described above for the different classes of mutations (see *Simulating fitness effects*), with results averaged across 10 independent replicates. In our results we display expected values of *B* as *B*_*q*_, where *q* is the central position in the chromosome. Thus, our *B* calculations represent the maximum effect of BGS across a chromosome, though when a simulated region is long enough to avoid significant edge effects (*L* ≥ 40000; *r* = 1 × 10^−8^), *B* for the central site can be considered to provide a general estimate of *B*_*q*_ across most sites (Table 1,Table S1; also see Figure 2b in Nordborg et al. 1996).

**Figure 1.**
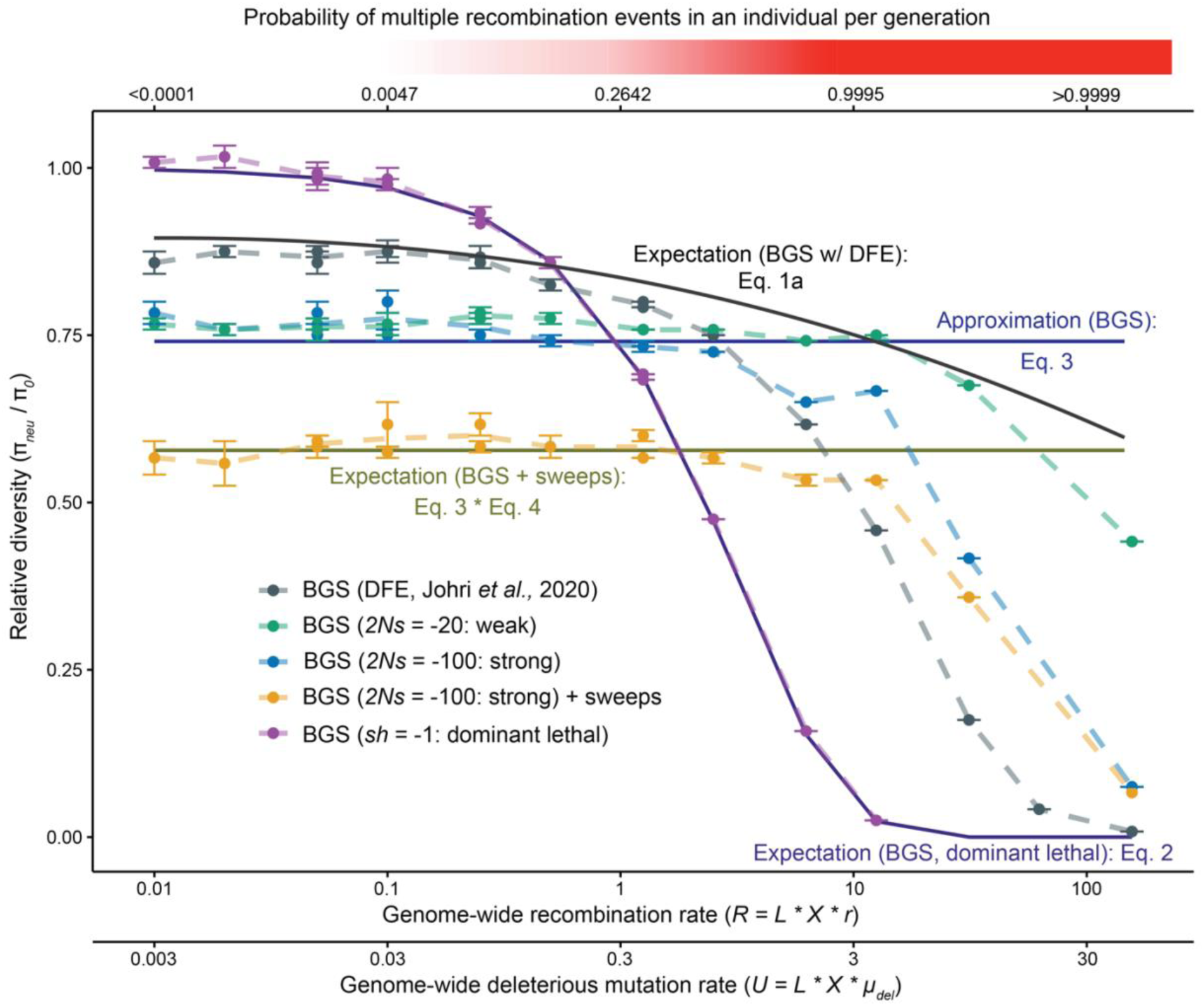
Observed nucleotide diversity at neutral sites in simulations with selection (π_*neu*_) relative to expected diversity under strict neutrality (π_0_ = 4*N*μ = 0.012) assuming a linear map distance, for simulations with varying combinations of length (*L*) and scaling factor (*X*), resulting in different genome-wide recombination rates (*R*) and deleterious mutation rates (*U*). Colors represent the different parameters of selection. Here, 50% of all mutations were selected. Points, connected by dashed lines represent the observations from simulations. Solid lines represent theoretical expectations calculated using Eq. 1a-4; the line using Eq. 1a is a LOESS regression line. The probability of multiple crossover events occurring in an individual per generation, *P*(> 1*CO*), corresponding to the simulated genome-wide recombination rate is indicated at the top. *N.B.*, only simulations with *L* ≥ 40 kb are shown. Standard error across 10 replicates is shown.

**Figure 2.**
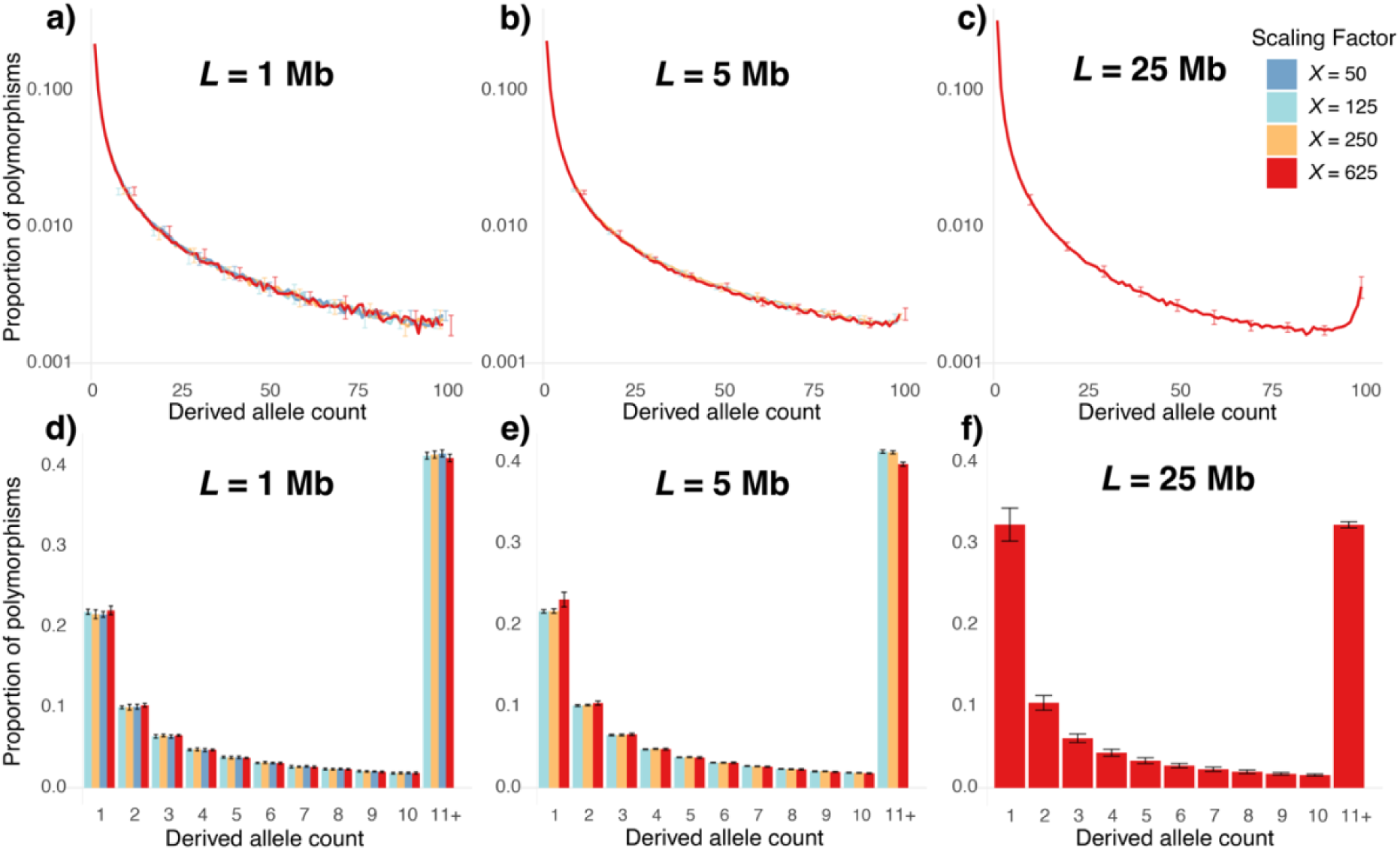
Site frequency spectra of derived neutral alleles where linked selected sites experienced strong purifying selection with constant selective effects (2*Ns* = −100) for regions of varying length (*L*). The different colours represent different scaling factors (*X*). The y-axis is presented on a log_10_scale for a-c. The final column in the bar plot represents the proportion of polymorphisms for all derived allele counts greater than 10 for d-f. Results are shown for a sample of 50 diploid individuals with standard error across 10 replicates. Results from simulations that were computationally infeasible were excluded.

When deleterious mutations are fully dominant (ℎ = 1) and lethal (*s* = −1), resulting in selected mutations never segregating in the population, the nucleotide diversity at neutral sites due to BGS relative to strict neutrality (*B*) depends solely on the rate at which the selected mutations are introduced (Nordborg et al. 1996), *i.e.*, the region-wide deleterious mutation rate per diploid individual per generation, *U*:

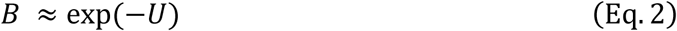

However, *B* is dependent on the region-wide recombination rate per individual per generation (*R*) when selection on new deleterious mutations is sufficiently mild so that deleterious alleles, while still typically purged from the population, may segregate and recombine. *B* can be approximated by the following expression for a central site without edge effects, provided *s*ℎ ≪ *R* (Hudson and Kaplan 1994; Charlesworth 2013):

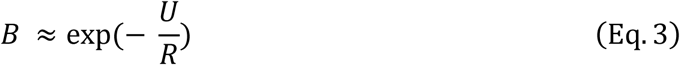

The assumptions used to obtain the above equation were met for our simulated parameters with *L* ≥ 40000 when simulating purifying selection of constant strength (2*Ns* = −100, −20; see *e.g.*, Table 1), but often were not when using the DFE that includes very strongly deleterious mutations. In Eq. 1-3, *B* reflects the relative nucleotide diversity due to background selection 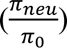 compared to that under neutrality in the absence of advantageous mutations. The nucleotide site diversity that takes into account the hitchhiking effects of selective sweeps can also be obtained from Stephan (1995):

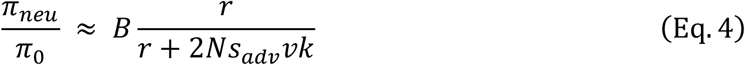

where 2*Ns*_*adv*_ is the beneficial population scaled selection coefficient, *v* is the rate of beneficial substitutions per site per generation which is equal to 4*Ns*_*adv*_μ_*adv*_ (μ_*adv*_ = 2μ × *f*_*pos*_ × *L*_*selected*_ being the region-wide rate at which new beneficial mutations are introduced per diploid individual per generation), and *k* is a constant approximately equal to 0.075. For simplicity Eq. 4 may be rewritten as the following:

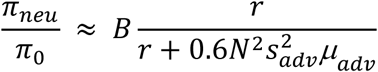

Finally, the mean expected reduction of nucleotide diversity as a function of genetic distance immediately following the completion of a selective sweep (ℎ), compared to before the beneficial mutation arose, was calculated using the following equation (Kim and Stephan 2000) for a site *i* bp from the beneficial fixed allele, where *c* is the probability of recombination between site *i* and the selected substitution, calculated linearly as *r* × *distance* (bp):

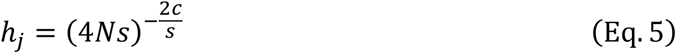

The mean expected diversity at a given neutral site j at time τ generations since fixation of the beneficial allele assuming no influence from other linked effects of selection (Kim and Stephan 2000) was approximated using:

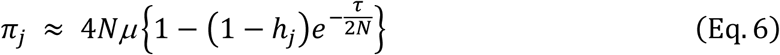

## Results

### Impacts of rescaling on diversity assuming recombination increases linearly with distance

To understand the effects of rescaling on neutral nucleotide diversity (π_*neu*_) in *D. melanogaster*-like genomes due to the linked effects of selection, we simulated all combinations of two parameters - the total length of the simulated region (*L* = 10 kb, 40 kb, 200 kb, 1 Mb, 5 Mb, 25 Mb), and the scaling factor (*X* = 25, 50, 125, 250, 625). We simulated a region with no genomic architecture, assuming that 50% of all new mutations experience selection. Note that *s*, the disadvantage or advantage of the mutant homozygote with respect to wildtype, was also scaled so that the population scaled selection coefficient (2*Ns*) had the same value or distribution in the rescaled simulations as in the unscaled population. While the simulated region-wide deleterious mutation rate (*U*) and crossover rate per individual per generation (*R*) varied across parameter combinations, they did so in proportion to the increases in *X* and *L*. If crossover interference is not simulated, which is the case in *SLiM*, an increase in *R* will lead to a proportional increase in the probability of multiple crossover events in each individual per generation, which we refer to as *P*(> 1 *CO*). As the number of multiple crossovers per individual increase, there is a decrease in the effective rate of recombination relative to the deleterious mutation rate, which might lead to an increase in linked effects of selection. Moreover, because the product of *X* and *L* determines both the probability of multiple crossover events in an individual as well as the number of deleterious mutations per individual, it must correlate with the effects of rescaling. As predicted, we observe a decrease in neutral diversity under different selection regimes (Figure 1) with an increase in *X* and *L*, as multiple crossovers become prevalent, compared to theoretical predictions that assume a linear genetic map (Eq. 1a, Eq. 3).

We observed that the nucleotide diversity at neutral sites with background selection relative to that under strict neutrality (*B*), generated by strong purifying selection with constant selective effects (2*Ns* = −100), overall matched our theoretical expectations (Eq. 1) quite well for regions of all length across all tested scaling factors (*X*). However, observed values of *B* decreased below the theoretical approximation that assumes a linear map (Eq. 3) when *L* ≥ 1 Mb and the scaling factor became larger than 250. In particular, the discrepancy between the theoretical expectation assuming linear approximation (Eq. 3) and results from simulations intensified as the product of the scaling factor and the length of the simulated region (*X* × *L*) increased *R* to where the probability of multiple crossovers in a single individual approached and exceeded 1 (observed 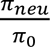 is 14% below the Eq. 3 approximation when *R* = 6.25; Figure 1). This largely occurred when simulated regions were large (*i.e*., 5 Mb or more) and when the scaling factor was as high as 625.

Similarly, when the strength of purifying selection was weaker (2*Ns* = −20), the observed 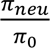 matched theoretical expectations assuming linear increase in the rate of recombination until *R* ≥ 31.25, at which point it decreased in a similar trend as that observed at lower *R* values for stronger selection parameters (Figure 1). When selective effects were drawn from a distribution of fitness effects (DFE; Johri et al. 2020), observed 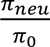 remained stable at around 0.87 in line with expectation (Eq. 1a) with increasing *X* × *L*, until *R* ≥ 1.25, at which point it began to decrease in a similar fashion to, albeit more precipitously than, the BGS simulations with constant selective effects (Figure 1).

Next, we introduced recurrent selective sweeps by adding a class of beneficial mutations under moderately strong selection 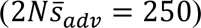, as estimated by Campos *et al*. (2017). These comprised 0.2% of selected mutations (*f*_*pos*_ = 0.002) while the remaining selected mutations experienced strong purifying selection of constant selective effects (2*Ns* = −100). We found that introducing recurrent selective sweeps reduced diversity in-line with theoretical expectations assuming a linear increase in the probability of recombination with distance (Eq. 4) for most combinations of scaling factor and length (Figure 1). However, diversity decreased below the theoretical expectation assuming a linear increase in recombination rate as multiple crossovers became prevalent when *R* ≥ 6.25 due to increasing *X* × *L*, similarly to what was observed when only BGS from strong purifying selection of constant selective effects was modelled (Figure 1).

When all selected mutations were dominant lethal with constant selective effects (*s* = −1; ℎ = 1, regardless of scaling factor), observed 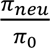 consistently matched theoretical expectations for non-segregating selected mutations (Eq. 2) at all parameter combinations (Figure 1; Table S3). Neutral simulations with no selection present exhibited diversity in-line with neutral expectations (π_0_ = 4*N*_*e*_μ = 0.012; Table S4) for all combinations of scaling factors and lengths.

The addition of gene conversion with BGS increased 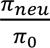 by up to 10%, as expected, though the same trend of decreasing diversity with increasing *X* × *L* was observed as in otherwise identical simulations experiencing strong purifying selection with constant selective effects (2*Ns* = −100) without gene conversion present (Table S5).

### Impacts of rescaling on diversity accounting for multiple crossovers

Because *SLiM* does not model crossover interference, a rescaling of the recombination rate per generation results in multiple crossovers per individual, that decrease the effective rate of recombination. We accounted for this effect theoretically by applying Haldane’s mapping function (Eq. 1b), which does not account for crossover interference and is therefore appropriate here. Theoretical expectations of neutral diversity with background selection for varying combinations of *X* and *L* are presented in Table 1. There are two main points to note. First, as *X* × *L* increases (and in turn *R* increases), there is a drastic expected decrease in neutral diversity due to BGS, showing a non-linear decrease in the coalescent *N*_e_. Secondly, while our theoretical predictions now get much closer to the neutral diversity observed in simulations (Table 1), they do not fully account for the observed decrease in 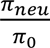.

When simulating BGS due to strong purifying selection with constant selective effects (2*Ns* = −100), *B* was overestimated using Eq. 1b by approximately 5% for *X* = 625; *L* = 25 Mb, and over two-fold for *X* = 625; *L* = 25 Mb, compared to simulated results (Table 1). Similar results were observed when selective sweeps were added (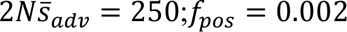; Table S6), and when simulating weaker purifying selection (2*Ns* = −20; Table S7). While the theoretical results accounting for multiple crossovers (Eq. 1b) also underestimated the effects of BGS from a DFE as multiple crossovers became prevalent, when *X* = 625; *L* = 25 Mb the simulations instead exhibited higher *B* than expected (Table S8). We therefore conclude that while multiple crossovers likely explain most of the increase in linked effects of selection, they do not fully explain biases observed due to rescaling. Note that, as expected, simulations of shorter regions were not particularly affected by multiple crossovers (Table 1). In addition, for the shortest simulations of *L* = 10 kb, the observed diversity was higher compared to *e.g. L* = 40 kb simulations which was well predicted by Eq. 1a (Table 1).

### Rescaling effects with different recombination-mutation ratios

Because rescaling results in a relative increase in both *R* and *U* (Figure 1; Table 1) in any simulated population, it is difficult to understand how rescaling effects would impact different organisms where the ratio of *U*/*R* may differ substantially from that in *Drosophila*. To assess the effects of a lower *U* relative to *R*, we modelled BGS due to strong purifying selection with constant selective effects (2*Ns* = −100) with the genome-wide deleterious mutation rate kept constant at a moderate level (*μ_scaled_* × *L* = *U* = 0.5) while other variables (*R* and *N*) scaled as expected with increasing *X*. While there was a slight decrease in the observed 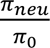 below theoretical expectations using the approximation (Eq. 3), there was not a drastic effect of increasing the value of *R* (as high as ∼150) as long as values of *U* were reasonably low (Table S9). Note however that increase in the linked effects of selection due to multiple crossovers only becomes relevant once *U* is sufficiently large. That is, only when there are multiple selected mutations on a chromosome that may recombine. Thus, it is difficult to successfully isolate the individual effects of *R* and *U*. This suggests that organisms with a lower ratio of *U* to *R* (*i.e.,* those that have genomes sparsely populated with functional regions) might preserve evolutionary dynamics at higher scaling factors.

We similarly assessed the effects of a higher *U* relative to *R* by keeping the genome-wide recombination rate constant at a moderate level (*r_scaled_* × *L* = *R* = 0.5) while *U* and *N* scaled as expected with increasing *X*. Again, for low values of *U*, the theoretical and observed values of 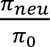 matched quite well. However, for *U* ≥ 2, observed values of 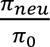 were much larger than the expected values using the approximation that assumes a linear increase in *R* (Eq. 3; Table S10). This suggests that the increased rate of deleterious mutation is likely playing an important role in generating the biases observed due to rescaling. Therefore, species that have a high ratio of *U* to *R*, i.e., those whose genomes are compact or have high rate of mutation, will likely be difficult to rescale accurately. Although note that when rescaling a population, both *R* and *U* will increase, thus understanding the effects of both remains crucial.

### Divergence

To assess the effects of *X* and *L* on divergence, the number of mutations reaching fixation (substitutions) from different classes (neutral, deleterious, beneficial where present) were tracked in the final 4*N* generations following a 10*N* generation burn-in, (Table 2, Table S11). Under all tested conditions, the rate of fixation of neutral mutations was constant, approximately equal to μ_*neu*_ per site per generation for each given *L* regardless of the *X* value (Table S11), as expected (Kimura 1968; Birky and Walsh 1988). Again, the number of fixations of deleterious mutations remained the same between simulated populations with scaling factors 25-250 and for regions smaller than 1 Mb in most scenarios (Table 2), suggesting that rescaling resulted in minimal biases in this parameter range. With much larger lengths and scaling factors, a dramatic increase in the number of deleterious fixations was observed, depending on the simulated parameters of selection (Table 2).

**Table 2.**
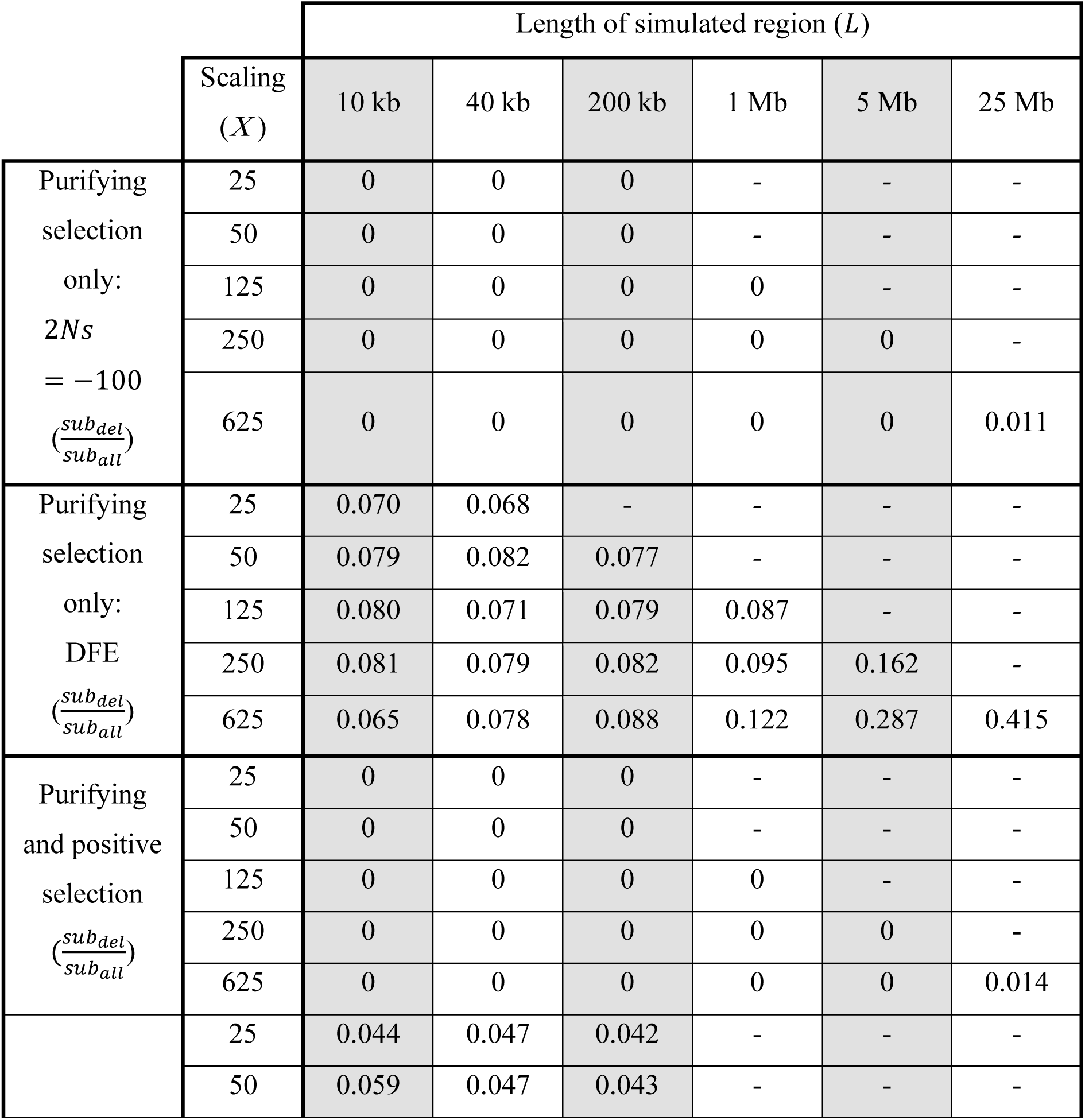
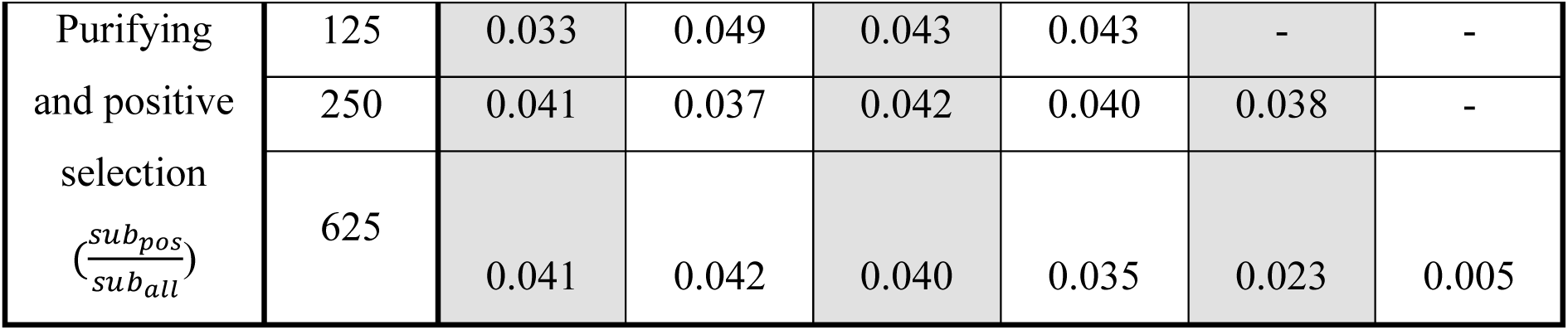
Proportion of all substitutions that were deleterious 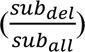 or beneficial 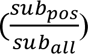 for simulations of varying length (*L*) and scaling factor (*X*). Half of all new mutations were selected. Three different evolutionary scenarios are shown: purifying selection simulated with constant selective effects (2*Ns* = −100), purifying selection with selective effects drawn from a DFE (Johri et al. 2020), and both purifying selection (constant 2*Ns* = −100) and positive selection modelled using parameters estimated by Campos *et al*. (2018; *f*_*pos*_ = 0.002; 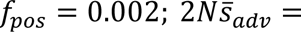 250), where 2*Ns* is the population-scaled strength of selection for selected alleles and *f*_*pos*_ is the proportion of selected mutations that were beneficial.

When simulating BGS due to strong purifying selection with constant selective effects (2*Ns* = −100), deleterious alleles did not fix under all parameter combinations, except *X* = 625; *L* = 25 Mb for which approximately 1.1% of substitutions were deleterious (Table 2). Simulating purifying selection with selective effects drawn from a DFE (Johri et al. 2020) that comprised 49% weakly deleterious mutations, resulted in approximately 8% of new substitutions being deleterious for most parameter combinations (Table 2). Note however that these substitutions are due to weakly deleterious mutations: the mean population-scaled selective strength of fixed deleterious alleles in these simulations was 2*Ns* = −2.58 across simulations with *L* = 200 Kb and the DFE. When *L* ≥ 1 Mb, increasing *X* did increase the proportion of substitutions which were deleterious, rising from approximately 8% at most parameters to 12.2% at *X* = 625; *L* = 1 Mb, and 41.5% at *X* = 625; *L* = 25 Mb, indicating that increasing proportions of weak and moderately deleterious mutations were reaching fixation (Table 2). However, it was only for the longest simulated region with the highest rescaling (*X* = 625; *L* = 25 Mb) that fixations of strongly deleterious mutations (2*Ns* ≤ −100) modelled by the DFE were observed (minimum one fixation per replicate), consistent with findings when simulating strong purifying selection with constant selective effects. Regions of length equal to or shorter than 200 Kb, with moderate rescaling factors (*X* < 625) provided highly accurate values of the proportion of deleterious substitutions.

The introduction of recurrent selective sweeps from beneficial mutations 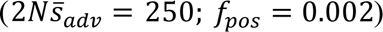 did not have a large effect on the fixation of strongly deleterious mutations (constant 2*Ns* = −100; Table 2), which is expected because the fixation probability of strongly deleterious mutations is negligible (Johri, Charlesworth, et al. 2021). In contrast, the proportion of substitutions that were beneficial decreased at high values of length × scaling factor, decreasing from approximately 4.2% when *L* = 200 Kb to 2.3% when *X* = 625; *L* = 5 Mb and 0.5% when *X* = 625; *L* = 25 Mb. In fact, the mean total number of beneficial fixations occurring in the final 4*N* generations was higher for *L* = 5 Mb; *X* = 250 (1165 beneficial substitutions) than *L* = 25 Mb; *X* = 625 (737 beneficial substitutions; Table S1), despite the latter having a five-fold increase in the number of beneficial mutations introduced. These results suggest that, while most rescaling parameters do not change fixation probabilities of new selected mutations, there is a significant and observable effect when the product of the length and the rescaling factor is high (e.g., when *X* ≥ 250 and *L* ≥ 1 Mb; Table 2; Table S11). There is an observed reduction in the efficacy of selection, resulting in an increase in the fixation of weakly deleterious mutations and a decrease in the fixation of beneficial mutations.

### SFS

The site frequency spectrum (SFS) of neutral sites was calculated with all sites under strict neutrality (Figure S1), and with BGS due to strong purifying selection with constant selective effects (2*Ns* = −100; Figure S2), BGS from purifying selection with selective effects drawn from a DFE (Johri et al. 2020; Figure S3), as well as BGS from purifying selection of constant strength (2*Ns* = −100) with recurrent selective sweeps 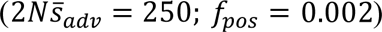. Under strict neutrality, there were no observable differences in the SFS across simulations with different combinations of *X* and *L* (Figure S4), apart from decreasing stochasticity with increasing *L*. Under all simulations with selection, no differences of note were observed for any tested scaling factors with *L* ≤ 1 Mb (Figure 2, Figure S2, Figure S3, Figure S4). However, when *X* = 625 and *L* ≥ 5 Mb, the proportion of variant sites at intermediate frequencies decreased, with increases in singletons and high frequency derived alleles (Figure 2, Figure S1, Figure S2, Figure S3). This change toward a “U-shaped” SFS was relatively mild for *X* = 625; *L* = 5 Mb (*R* = 31.25; *U* = 9.375), though it became severe when *X* = 625; *L* = 25 Mb (*R* = 156.25; *U* = 46.875), especially when BGS was modelled with a DFE (Johri et al. 2020; Figure S2) with a large proportion of mildly deleterious mutations. Again, note that even with simulations of regions as large as 5 Mb and a scaling factor of 250, with a DFE relevant to *D. melanogaster* populations, there were no observed effects of rescaling in the SFS.

### Linkage disequilibrium

The decay of pairwise linkage disequilibrium (LD) between neutral sites was consistent across tested rescaling parameters under strict neutrality (Figure S5). It was similarly unaffected by rescaling for much of the parameter space, including when simulations were carried out using 5 Mb regions with *X* = 250. Similarly to the SFS, LD decay became perturbed only when the length of the simulated region and the scaling factor were large (*i.e.,* at *X* = 625; *L* = 5 Mb and *X* = 625; *L* = 25 Mb; Figure 3, S6, S7). When simulating BGS generated by strong purifying selection with constant selective effects (2*Ns* = −100), LD did not change with increasing *X* and *L* until *X* = 625; *L* = 5 Mb, when there was a slight flattening of LD decay with increasing distance between pairs of neutral alleles (Figure 3). This effect became noticeable only at *X* = 625; *L* = 5 Mb, at which point mean *r*^2^ between sites 1 kb apart increased from approximately 0.0210 to 0.0245, compared to what was seen with milder rescaling parameters (Figure 3). Modelling BGS instead with a DFE (Johri et al. 2020) led to more extreme effects, for *X* = 625; *L* = 5 Mb mean *r*^2^ between sites 1 kb apart increased further to 0.426 (Figure S6), with substantial changes in the shape of the decay rate. The addition of selective sweeps to simulations with BGS from strong purifying selection with constant selective effects (2*Ns* = −100) did not appear to change the effects of rescaling on LD any more than with BGS alone (Figure S7).

**Figure 3.**
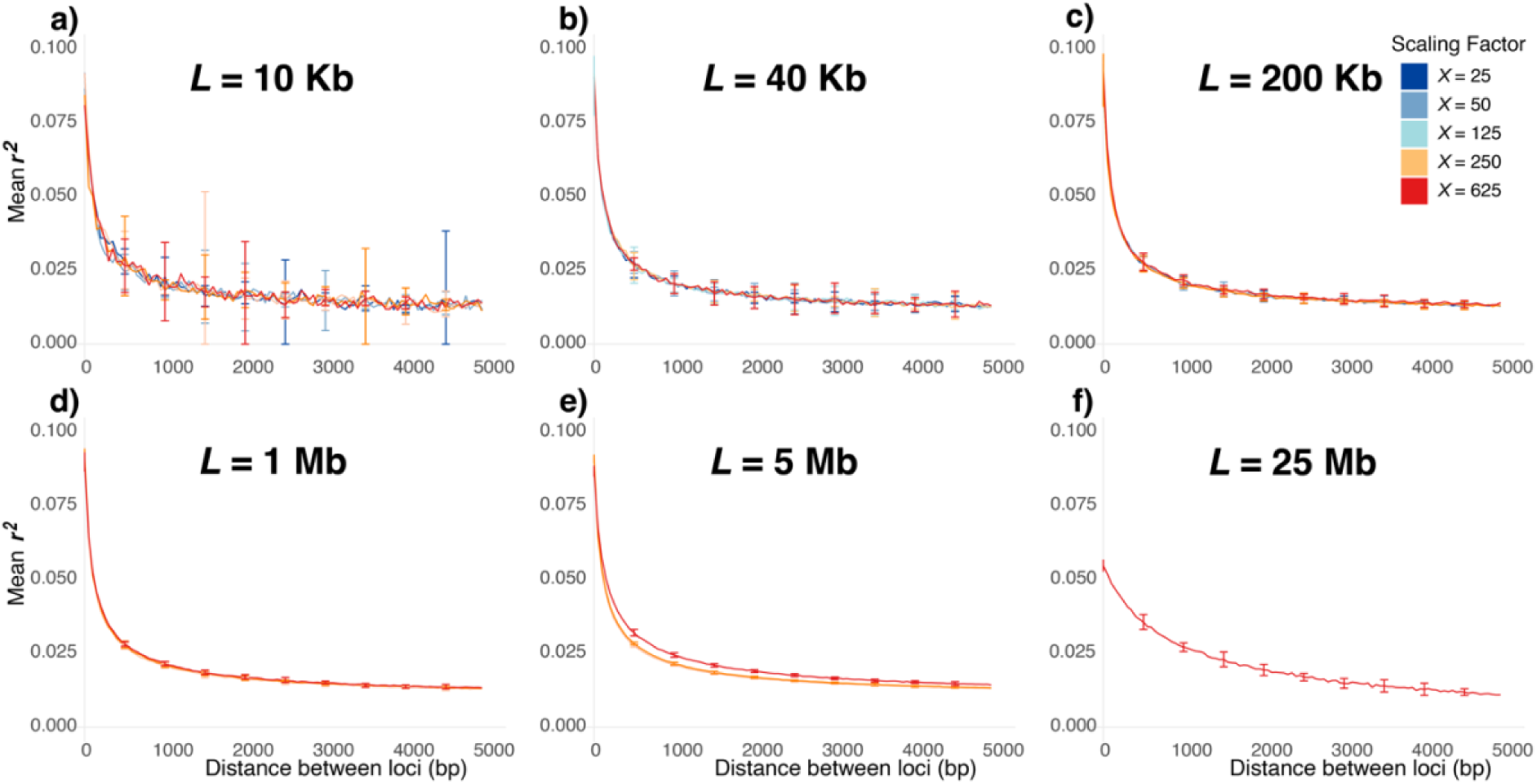
The decay of linkage disequilibrium (measured by *r*^2^) as a function of distance between all pairs of alleles from simulations experiencing strong purifying selection with constant selective effects (2*Ns* = −100) for regions of varying length (*L*; a-f). Here, 50% of all mutations were selected. The different coloured lines represent results from simulations with different scaling factors (*X*), excluding parameters beyond computational constraints. Results are from 50 diploid samples and standard error across 10 replicates is shown.

To test whether the flattening of LD decay for neutral sites in highly rescaled simulations experiencing BGS was due simply to a reduction in effective population size, we simulated a strictly neutral region with an equivalent population size for comparison. We calculated the effective population size in simulations experiencing BGS from strong purifying selection with constant selective effects (2*Ns* = −100) with the highest length × scaling (*X* = 625; *L* = 25 Mb): given 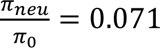 (Table 1), we reduced *N_scaled_* to *N_scaled_* × 0.071 and simulated using otherwise identical parameters, though under strict neutrality. We found neutral simulations with reduced *N* exhibited far higher LD (Figure S8), indicating the observed flattening of the LD curve in very high *X* × *L* simulations experiencing selection could not be explained only by the reduction in *N*_*e*_ due to selection.

### Offspring distribution and fitness dispersion

To test for progeny skew in populations experiencing selection we examined the distribution of offspring counts per individual, *i.e.*, the number of progeny (in haplotypes) each haplotype in the parental population contributes to the following generation, for the final 100 generations in select simulations. We observed a substantial increase in progeny skew for highly rescaled simulations modelling strong purifying selection of constant strength (*X* = 625; 2*Ns* = −100) as the length of the simulated region increased from *L* = 200 kb to *L* = 25 Mb: the maximum offspring count observed in the final 100 generations for a haplotype increased fivefold for the longer simulated region (13 offspring to 72 offspring) and the proportion of haplotypes in the parental population with no offspring increased from 14% to 36% (Figure 4). This effect was more extreme when modelling purifying selection with a DFE (Johri et al. 2020), due to the increase in dispersion of fitness between individuals (see below), where the maximum offspring count observed for a haplotype increased two orders of magnitude with the higher length (*X* = 625; *L* = 200 kb to *X* = 625; *L* = 25 Mb). As a result, when *X* = 625; *L* = 25 Mb, a single haplotype was observed to have 1420 progeny of the total 3200 (across 1000 total generations tested), providing 44% of the genetic information to the following generation. In addition, the proportion of haplotypes in the parental population with no offspring increased from 14% to 66% (Figure 4). We found that the variance in the offspring counts became non-Poisson as the observed variance far exceeded the mean in *X* = 625; *L* = 25 Mb simulations experiencing selection (Figure 4), violating Wright-Fisher assumptions. Note that while an increase in the proportion of haplotypes in the parental populations that contribute to no offspring in the next generation are expected to increase with an increase in the extent of BGS (by definition), the entire offspring distribution (excluding the haplotypes contributing no offspring) was significantly different in the highly rescaled simulations (Figure 4). Thus, the observed progeny skew was unlikely to be a result of simple rescaling of the effective population size due to BGS.

**Figure 4.**
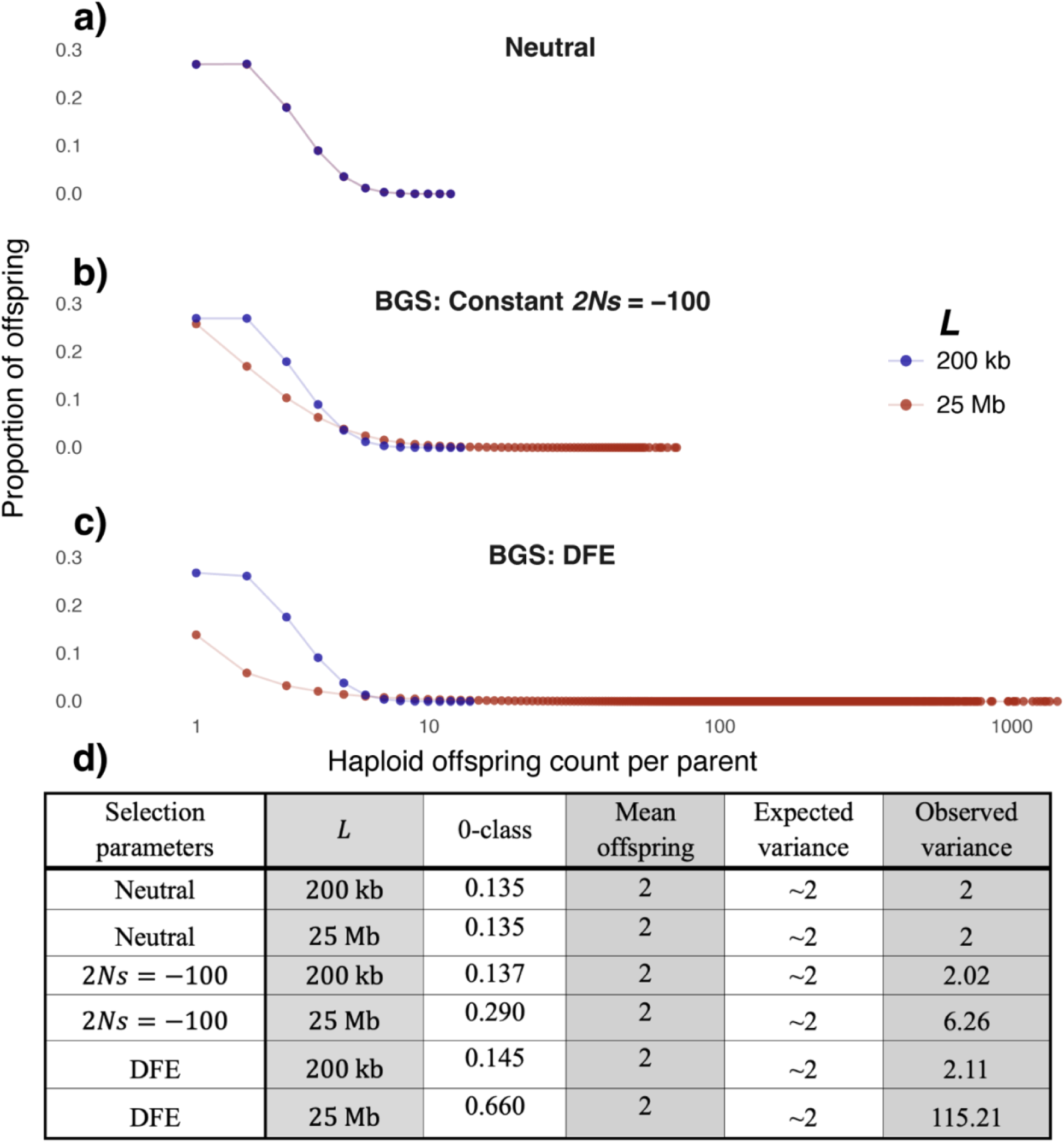
Offspring distributions recorded in the final 100 generations of simulations (following burn-in) under (a) strict neutrality and (b-c) those experiencing purifying selection (where 50% of all mutations were selected). Here we show cases where the scaling factor (*X*) is large, with increasing length of the simulated region (*L*). In all simulations, *X* = 625 with 1600 diploid individuals (3200 haplotypes), which is the total number of progeny across all parents per generation. (d) The table shows the corresponding proportions of “0-class” individuals, *i.e.,* individuals in the parental generation that did not have offspring, the mean offspring count (in haplotypes) per parental individual (“mean offspring”), the expected variance in offspring count per individual under Wright-Fisher dynamics, and the observed variance in offspring count per individual from simulations. Results are shown from 10 replicates.

We also tracked the mean and variance of fitness per individual in the final 100 generations for the same simulations and found drastically increased dispersion in fitness in populations experiencing progeny skew due to high *X* × *L* (Table S12). In simulations modelling the DFE, the coefficient of variation of per-individual fitness (*SD/mean*) increased from 0.02 when *X* = 125; *L* = 200 Kb to 0.08 when *X* = 625; *L* = 200 Kb, and up to 1.37 when increasing the length to *L* = 25 Mb (*X* = 625). Similar, though more extreme results were observed when modelling the DFE, where the coefficient of variation reached 10.06 when *X* = 625; *L* = 25 Mb (Table S12). This indicates that the increased dispersion in fitness associated with progeny skew is increasing as a function of scaling factor and length. In addition, the mean fitness of very long, highly rescaled simulations experiencing selection (*X* = 625; *L* = 25 Mb) was extremely low, ≤ 8.3 × 10^−9^, presumably from the fixation of deleterious mutations, whereas it was above 0.6 in all tested 200 kb simulations.

### Local linked effects of recurrent selection

To test the local impacts of rescaling on the linked effects of selection, in particular the effects on neutral sites adjacent to a large, conserved region such as an exon, we calculated summary statistics for sites in a strictly neutral region as a function of distance (bp) from a 10 kb region under selection simulated under different scaling factors (*X*). When the selected region experienced only purifying selection with selective effects modelled by the DFE from Johri *et al.,* 2020 (where 75% of sites experienced selection), there were minimal effects of BGS on diversity with increasing distance from the selected region, and this was consistent across scaling factors (Figure 5). Local diversity was more substantially reduced when recurrent selective sweeps 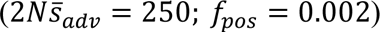 were modelled with purifying selection (as above with DFE; Johri et al. 2020), so that neutral sites in the 250 bp immediately adjacent to the selected block had approximately 10% lower diversity than neutral sites 10 kb from the selected block (Figure 5); however, there were no substantial differences observed across scaling factors.

**Figure 5.**
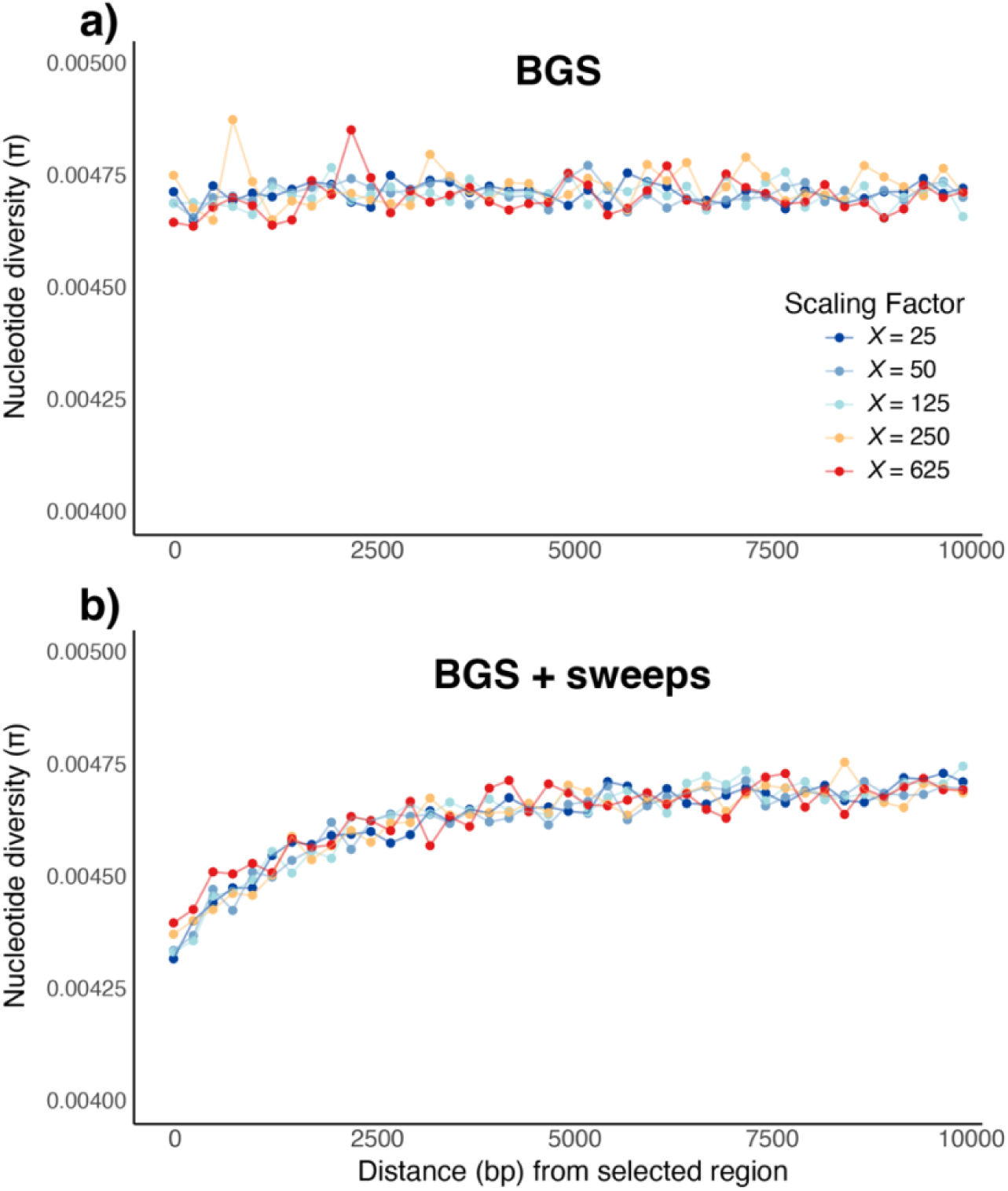
The recovery of nucleotide site diversity (π) observed for sites in a strictly neutral region as a function of distance from an adjacent 10 kb block within which all sites experienced (a) purifying selection (DFE from Johri *et al.,* 2020), or (b) purifying and recurrent positive selection 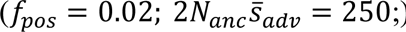, from simulations with different scaling factors (*X*). Results have been averaged across 10000 replicates.

We also calculated alternative summary statistics commonly used to test for signatures of selection, Tajima’s *D* (Tajima 1989) and Fay and Wu’s *H* (Fay and Wu 2000), as a function of distance from the 10 kb selected block, though no clear trends emerged across scaling factors under either BGS only or BGS with sweeps (Figure S9, S10), indicating they are not likely to be severely affected by rescaling.

### Recovery of neutral diversity from a selective sweep

To test the impacts of rescaling on the local effects of selective sweeps, we simulated a 5 kb neutral region and introduced a single beneficial mutation of constant selective strength which swept to fixation, observing diversity of neutral sites as a function of distance from the selected site at specified time intervals, comparing with theoretical expectations when possible (Eq. 6; Figure 6). When the introduced beneficial mutation was moderately strong (2*Ns*_*adv*_ = 100), no substantial differences were observed between scaling factors, with all parameters producing reduced mean diversity at the time of fixation, consistent with the theoretically predicted recovery of mean diversity post-fixation (Figure 6). When the introduced selected mutation was ten-fold more advantageous (2*Ns*_*adv*_ = 1000) the observed mean diversity of sites with distance greater than approximately 400 bp from the selected site was slightly higher than theoretical predictions for approximately 3*N* generations post-fixation (Figure S11). As expected, this effect did not substantially differ across the tested scaling factors (Figure S11).

**Figure 6.**
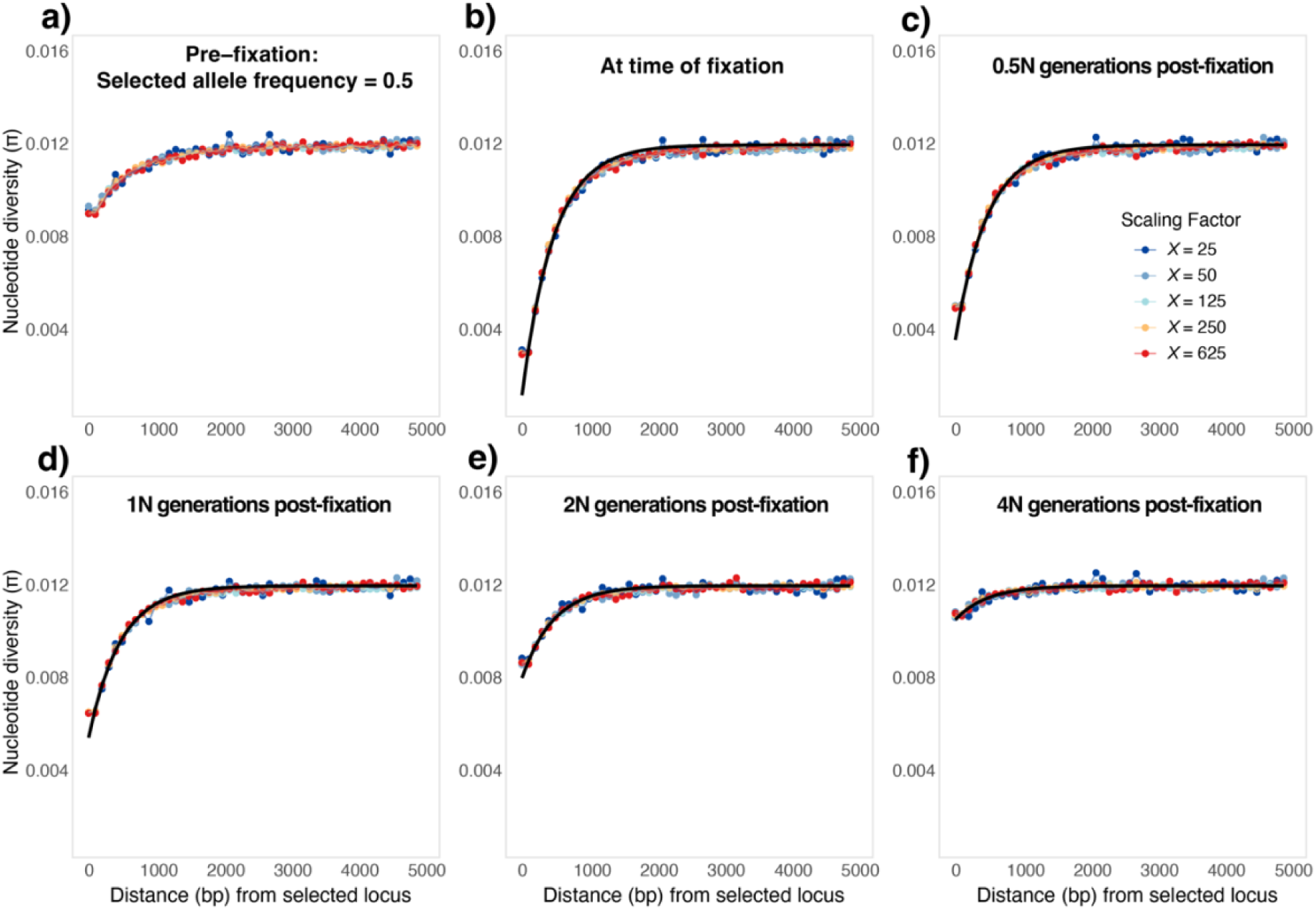
The recovery of nucleotide site diversity (π) observed for sites in a strictly neutral region as a function of distance from the fixation of a single moderately strong beneficial mutation (2*Ns*_*adv*_ = 100). Results are shown for different scaling factors (*X*) and for varying times during and post-fixation (as indicated on top of each plot). Here, results are aggregated across a minimum of 1792 replicates (coloured dots). The black line represents the expected diversity post-fixation calculated using Eq. 6. For an interactive plot showing data from all measured time points, see https://jacobimarsh.github.io/SweepRecov/.

## Discussion

We have here demonstrated that, for a large parameter space relevant to *Drosophila*-like populations, rescaling effects are negligible or minor. They become severe when the rescaling factor becomes very large (e.g. *X* = 625) and when a large number of linked selected sites are simulated (usually larger than 1 Mb chromosomal regions). While this is good news for many researchers interested in simulating shorter regions in *D. melanogaster*, it seems that simulating full-length chromosomes accurately is not currently possible for populations that have large effective sizes. It is, therefore, still useful to understand which factors are responsible for the observed rescaling effects and how to perform simulations in the parameter space unaffected by biases introduced by rescaling. Below we discuss the likely factors responsible for such biases.

### Diffusion limits

The standard diffusion approximations we use that calculate expected diversity rely on underlying assumptions of μ → 0, *s* → 0, and large population size limit (*N* → ∞) such that the combined quantities, *N*μ → *finite* (Norman 1975; Wakeley 2005), *rL* → 0, and *NrL* → *finite* (Ewens 2004; Barton and Otto 2005; Innan and Sakamoto 2021) describe the probability density function of the allele frequency trajectories. Since the diffusion process is a continuous time Markov process, the transition from discrete to continuous variables is carried out by ignoring higher order terms of μ, *s*, and *r* such that the terms μ^2^, *s*^2^, *r*^2^, and 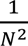 are ignored. Thus under the diffusion limit, a number of population-genetic quantities of interest scale linearly with *N*μ, *Ns*, and *NrL*, providing the basis for laws of rescaling. In rescaled simulations of relatively small genomic regions, if the above-mentioned theoretical assumptions are met, it is unlikely that any biases will arise.

Although the condition, μ*X* ≪ 1, is usually easy to satisfy in rescaled populations and also holds in the current study, *rLX* ≪ 1 is not satisfied when simulating extremely long regions with high rescaling, *i.e*., when *L* × *X* reach the point where multiple crossovers become prevalent, and crossover probability is no longer linearly correlated with distance (*i.e.*, Haldane’s mapping function is necessary; Haldane 1919). Similarly, when a very small population size is simulated during a bottleneck event, 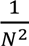 may no longer be negligible and might lead to deviations from observations in an unscaled population. Finally, when simulating strong selection, *s*ℎ*X* can be close to 1, thus potentially violating assumptions of diffusion theory. When simulating BGS with fixed selective effects, |*s*ℎ*X* | reached only 0.156 (when *X* = 625) for simulations with 2*Ns* = −100, although it approached 0.5 for the strongly deleterious class when simulating the DFE. When simulating recurrent selective sweeps (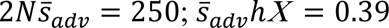 when *X* = 625) our theoretical expectations matched observed diversity well for short regions (Table S6). While we did not observe any substantial effects of rescaling when simulating highly beneficial mutations with *s*ℎ*X* approaching 1 or being above 1, researchers should be careful when simulating specific values of “*s*”, especially if their parameters lead to violating the diffusion approximations. Thus, these are additional factors that researchers should keep in mind while designing their simulations.

### Effects of simulating short regions: increased diversity and stochasticity

When simulating short regions (*L* = 10 kb), high stochasticity between replicates was present regardless of the scaling factor (Table 1), as we only tested 10 replicates of each condition. High stochasticity between replicates from simulations with low *L* may be explained by the reduction in information (sites) analysed and may be counteracted by increasing the number of replicates (Dabi and Schrider 2024; Ferrari et al. 2024). We did not observe increasing *X* having a consistent impact on stochasticity for diversity, divergence, LD, SFS, and the linked effects of selection, for regions of small to moderate length (*L* ≤ 200 kb).

The effects of BGS decreased for *L* = 10 kb compared to longer regions, regardless of scaling factor, as expected from *B* given by Eq. 1b (Table 1), though deviating from the approximation of *B* from Eq. 3. This is due to the fact that most of the simulated region consists of “edges”, which reduces the linked effects of selection (Charlesworth 2013).

### Effects of high rescaling: multiple crossovers

Multiple crossovers during the generation of each individual become increasingly common with increased rescaling and the length of simulated region (*R*_*scaled*_ = *X* × *L* × *r*_*unscaled*_), especially as *SLiM* assumes no crossover interference. Multiple crossovers significantly change the dynamics of genetic inheritance when compared to a population where only single crossovers per individual per generation are likely to occur (Haldane 1919). For example, when two crossover events occur, the distal ends of the resultant (double) recombined chromosome will be fully linked as the second recombination event reverses the effects of the first for downstream loci (Figure 7a). Thus, crossover probability between two points far away is less correlated with the physical distance between them in a highly rescaled population experiencing multiple crossovers than it would be across the same physical distance in an unscaled population.

**Figure 7.**
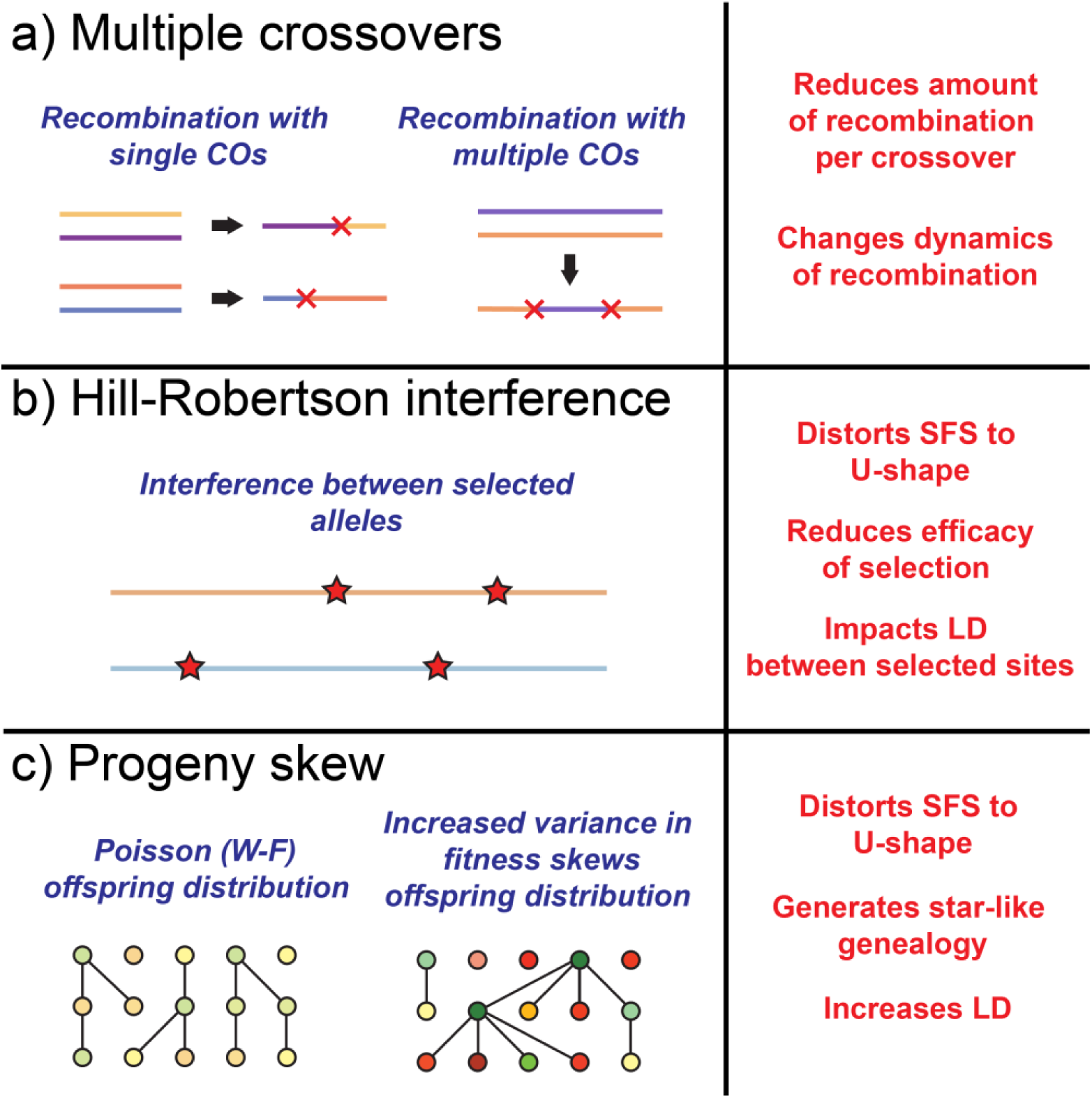
Schematic diagrams of the effects of excessive rescaling (left) which change population genetic dynamics and statistics (right). In section a), each coloured line represents a haplotype, with crossover breakpoints indicated as crosses; b) each coloured line represents a haplotype, with selected mutations indicated as stars; c) a simplified genealogy where each node represents a haploid individual and connections between parent-offspring, colours indicate fitness effects (red for low fitness, green for high fitness).

When a single crossover occurs in the centre of the chromosome, the maximal rate of genetic exchange, *i.e.,* the maximum proportion of the genome that recombines between the two parental haplotypes, is 50%. When multiple crossovers occur, on average the rate of genetic exchange will increase toward 50% as the chances of only recombining at the ends of the chromosome decrease, though each crossover beyond the first will result in a diminishing average increase in the rate of genetic exchange as it trends toward 50% (Haldane 1919). As a result, the effective rate of genetic information recombining per generation will be reduced with large increases to length × scaling factor (*L* and *X*) which increases the rate of multiple crossovers. This effect of multiple crossovers, i.e., reduced effective recombination genome-wide, might not significantly impact summary statistics such as nucleotide diversity and the SFS when most alleles are neutral. However, when purifying selection is present they will exhibit increased effects of BGS as the probability of alleles recombining onto new higher relative fitness genetic backgrounds decreases.

While we predict that multiple crossovers may significantly impact chromosome-wide dynamics of recombination, local dynamics are expected to be preserved. For our highest scaling factor, *X* = 625, the mean distance between two independent recombination events is 160 kb = 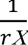. However, in our simulations, the decay of mean *r*^2^ with distance plateaus beyond approximately 4 kb (Figure 3). Thus, LD between two sites with physical distance 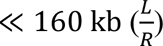 is expected to be unaffected by multiple crossovers while *X* ≤ 625, regardless of the total length of the region simulated. Therefore, local LD, including the local linked effects of selection (*e.g.,* a selective sweep), is not expected to be impacted by multiple crossovers occurring in a single generation, unless the sweep is very strong and extremely recent.

Finally, it should be noted that while multiple crossovers do take place in *Drosophila* populations (Sturtevant 1913; Aggarwal et al. 2015), the co-incidence rate of secondary crossovers is strongly influenced by proximity to other chiasmata on the same chromosome – a phenomenon known as crossover interference (Muller 1916; Berchowitz and Copenhaver 2010; Otto and Payseur 2019). In *Drosophila* species, crossover interference is typically strong relative to other species (Aggarwal et al. 2015), in *D. yakuba* multiple crossovers occur for approximately 10% of each chromosome per offspring (Pettie et al. 2022). Other species can exhibit very different crossover rates, with widely varying strengths of crossover interference (Otto and Payseur 2019). In *Caenorhabditis elegans*, crossover interference is almost complete (Barnes et al. 1995). As a unique example, upwards of 10 crossovers per chromosome is common in the fission yeast *Schizosaccaromyces pombe* for which typical crossover interference is not present (Munz 1994; Cromie and Smith 2008). To effectively model such a population it may in fact be most appropriate to simulate a matching number of multiple crossovers per chromosome.

Crossover interference is typically not modelled in forward-genetic simulators such as *SLiM* (Haller and Messer 2023) and *fwdpp/fwdpy* (Thornton 2014; Thornton 2019); to model crossover interference would require the implementation of mapping functions (*e.g.,* Zhao and Speed 1996). Therefore, even for populations with high rates of multiple crossovers, realistic dynamics of crossover interference for most species are not effectively simulated in highly rescaled simulations when multiple crossovers occur. Finally, it’s essential to note that even realistic crossover interference was modelled in simulations, a population that is rescaled will be more severely impacted by crossover interference effects due to the higher crossover rate compared to the unscaled population. In other words, there would be an upper limit to the scaling factor that would allow the rescaled population to have the same effective recombination rate as the unscaled population.

### Effects of high rescaling: interference and progeny skew

Accounting for multiple crossovers using a Haldane map (Eq. 1b) enabled more accurate predictions of diversity in highly rescaled populations, though there was still a mismatch between analytical and simulated results (Table 1, S6, S7, S8), indicating additional sources of bias. Hill-Robertson interference among loci under selection (HRI; Hill and Robertson 1966; Felsenstein 1974; Otto 2021), can occur as a result of very high rescaling, as first pointed out by Comeron and Kreitman (2002). In highly rescaled simulations (*e.g., X* = 625; *L* ≥ 5 Mb) an increase in HRI occurs from (a) the increase in the genome-wide deleterious mutation rate per individual per generation (U; see Uricchio and Hernandez 2014 for related analysis and discussion with rescaled simulations modelling positive selection), and (b) a corresponding reduction in the effective recombination rate genome-wide from multiple crossovers. A decrease in the effective rate of recombination not only increases the extent of background selection, as observed in this study (Figure 1 and Table 1), but also lowers the probability of less-deleterious combinations of alleles recombining onto the same haplotype, reducing the ability of the population to purge deleterious alleles (Comeron and Kreitman 2002; Kaiser and Charlesworth 2009). Thus, HRI, including the effects of background selection and selective sweeps (Charlesworth and Jensen 2021), diminishes the efficacy of selection (Figure 7b).

HRI can cause perturbations to several population genetic summary statistics, including the SFS, divergence, and LD (Comeron et al. 1999; McVean and Charlesworth 2000; Comeron and Kreitman 2002; Garcia and Lohmueller 2021; Becher and Charlesworth 2025). Populations experiencing HRI have been reported to exhibit a non-monotonic SFS with a characteristic U-shape (Good et al. 2014; Zeng and Corcoran 2015; Cvijović et al. 2018), suggesting that increased HRI caused by very high length × scaling factor may be partially responsible for the observed U-shaped SFS in simulated populations experiencing BGS (Figure 2). Moreover, an increased rate of fixation of deleterious mutations in highly rescaled populations also suggests the role of HRI (Comeron and Kreitman 2002; Table 2). In highly rescaled simulations with selection we observed progeny skew in the offspring distribution which was associated with increased dispersion of fitness between individuals relative to mean fitness across the population (Table S12; Figure 4, Figure 7c). This severe progeny skew violates Wright-Fisher dynamics which assume that offspring counts are Poisson-distributed with a variance equal to the mean.

When this is violated and the variance is much larger, it introduces significant biases to population genetics summary statistics (Irwin et al. 2016). An effect of severe progeny skew is the increased incidence of multiple merger events, where several lineages coalesce simultaneously, thus violating the Kingman coalescent assumed in Wright-Fisher processes (Irwin et al. 2016; Matuszewski et al. 2018), multiple merger events are also typically associated with a U-shaped SFS (Eldon and Wakeley 2006; Blath et al. 2016; Freund et al. 2023). Thus, it is possible that the U-shape of the SFS for neutral sites (observed in the presence of background selection alone) was also caused by a dramatic increase in progeny skew toward a sweepstake reproduction-like regime, where highly skewed distributions of offspring numbers per individual result in few parental haplotypes contributing genetic information disproportionally to the following generation (Williams 1975).

To better understand how progeny skew occurs in highly rescaled simulations and how it impacts populations, we must consider the effects of high variance in fitness between individuals introduced by a very high deleterious mutation rate per individual per generation (*U*) relative to the effective rate of recombination. When high variance in fitness is introduced from generation to generation, such as for the parameters *X* = 625; *L* ≥ 5 Mb where *U* ≥ 9.375, the “least-deleterious” genomic background will become substantially more fit than the average fitness across the population, resulting in a sweep-like effect which decreases the proportion of intermediate frequency alleles (see Williams 1975; Durrett and Schweinsberg 2005; Árnason et al. 2023 for related discussions in populations experiencing sweepstake reproduction due to beneficial mutations). This sweep-like effect is likely responsible for the fixation of strongly deleterious alleles observed in highly rescaled simulations when they are linked to a “least-deleterious” genomic background, rather than a non-deleterious background which is not common given the value of *U* (Table 1). In a finite population where population size is dependent on the fitness of the previous generation, this level of recurrent fixation of deleterious mutations would induce a mutational meltdown due to Muller’s ratchet (Muller 1964) where the population rapidly enters an extinction vortex due to the inability of recombination to effectively purge deleterious mutations in successive generations whilst new deleterious mutations arise (Lynch and Gabriel 1990; Gabriel et al. 1993; Lynch et al. 1993; Lynch et al. 1995). In this regime, the fixation probability of new beneficial mutations approaches zero under sufficiently high *U*, as the fitness advantage of a new advantageous allele is overshadowed by the relative decrease in fitness caused by the introduction of vast numbers of new deleterious mutations (Pénisson et al. 2017). Multiple-merger events can increase the efficacy of positive selection (Der et al. 2011). However, when the selective strength of beneficial mutations is not extremely strong and the proportion of selected mutations that are beneficial is very low, as in our simulations 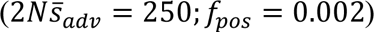, we expect this selective effect of the beneficial mutation to be trivial compared to the impact of multitudinous new deleterious mutations when determining relative fitness across individuals. As a result, the overshadowing effect of abundant deleterious mutations (Pénisson et al. 2017) is a likely explanation for the substantial decrease in fixation probability of beneficial mutations in highly rescaled simulations (Table 2). Note that while Muller’s ratchet generally occurs in a non-recombining population, in our case it is due to the high ratio of *U* relative to *R*.

Finally, skewed offspring distributions have also been shown to increase LD when the offspring of a single individual replaces an intermediate fraction of the population, as seen in simulations with extremely high length × scaling (Figure 4; Eldon and Wakeley 2008), which best explains the patterns of LD decay observed in highly rescaled populations.

It is necessary to reiterate, that while the effects of high rescaling related to HRI and progeny skew may profoundly affect population genetic dynamics, we only observed such effects in simulations of large regions with a very high scaling factor (*X* = 625; *L* ≥ 5 Mb with strong purifying selection of constant strength, *X* = 625; *L* ≥ 1 Mb when simulating a DFE), likely due to a very high deleterious mutation rate (*U*).

### Comparison with recent rescaling studies

Our results provide an opportunity to compare and reflect on findings reported by recent studies assessing the impact of rescaling on forward population genetic simulations. Dabi and Schrider (2024) observed minimal effects of rescaling for *Drosophila*-like simulations of 10 kb segments with scaling factors ranging from *X* = 20 to *X* = 200. They did however find stronger effects of rescaling when extremely strong recurrent selective sweeps are present (2*Ns* > 10000), though in these simulations *s*ℎ*X* = 1 when *X* = 200, so that the diffusion approximation with respect to selection is not expected to hold (see ‘Diffusion limits’ above). In light of our results, it appears that they did not observe more severe effects of rescaling with most *Drosophila*-like parameters due to the limited length of the simulated region (*L* = 10 Kb) which keeps values of the parameters *U* and *R* (which are dependent on *L* × *X*) low enough to avoid introducing multiple crossovers, HRI and progeny skew (Figure 7). However, beyond *Drosophila*, the authors also tested larger regions (*L* = 25 Mb) with human-like parameters, observing substantial decreases in the fixation probability of beneficial mutations with increasing scaling factor (*X* = 2 to *X* = 20), while the fixation probability of deleterious mutations increased, LD across all sites increased, and the proportion of singleton alleles rose (under the “no-beneficials” model, see Figure S3b, S5b in Dabi and Schrider (2024). While the details of our study and theirs make direct comparisons difficult (*e.g.,* differences in *N* and selective effects), their results match well with what may be expected from our findings, as at their highest scaling factor, *R* = 6, indicating that multiple crossovers would occur in approximately 98% of new individuals (Dabi and Schrider 2024). Moreover, their highest simulated mutation rate was *U* = 6 which was probably sufficient to produce HRI and/or progeny skew effects. This was approximately the region-wide mutation rate with our parameters that produce a DFE needed to increase LD (Figure S6) and distort the SFS toward a U-shape with high proportions of singletons (Figure S3) while decreasing the fixation probability of beneficial mutations (Table 2) and increasing the fixation probability of deleterious mutations (Table 2). It should be noted that the proportion of singletons in the SFS did not increase with increased scaling factor when Dabi and Schrider (2024) simulated these parameters with the addition of beneficial mutations. Nevertheless, the shared biases due to rescaling seen in our *Drosophila-*like simulations and their human-like simulations under similar *U* and *R* highlight the role of these factors in producing effects that alter population genetic dynamics in parameter spaces relevant to a wide array of species.

Ferrari *et al*. (2024) conducted a wide array of forward-in-time simulations with selection, with human-like and *Drosophila*-like parameters, testing some combinations of genome length (*L*) and scaling factor (*X*); they observed several effects consistent with our findings. Under *Drosophila*-like parameters, they observed a reduction in diversity when simulating long regions with high rescaling (*L* ≥ 1 Mb; *X* ≥ 500), similar to our results presented in Figure 1, as well as a reduction in intermediate allele frequencies when simulating very long regions with high rescaling (*L* ≥ 10 Mb; *X* ≥ 500), similar to our results presented in Figure 2. The authors conclude that the decrease in diversity in long simulated regions with high rescaling is due to increased effects of selection (Ferrari et al. 2024). We would, however, conclude that the decrease in recombination rate induced by multiple crossovers, as well as the increase in the number of deleterious mutations per genome leading to HRI, and progeny skew, are causing the biases due to rescaling (as discussed above), particularly because they simulated *R* values up to 100 and *U* values up to 62.18 for *D. melanogaster*-like populations (Ferrari et al. 2024).

Adrion *et al*. (2020) found that forward-in-time population genetic simulations in *SLiM* modelling human-like populations under strict neutrality (no selection) exhibited reduced LD and diversity under high rescaling (*L* = 51 Mb; *X* = 50); however, in our results modelling strict neutrality we did not observe directional biases associated with any summary statistics.

Given that their demographic model includes extreme bottlenecks (Kamm et al. 2020), it is possible that their reported effects could be due to population sizes temporarily dropping to extremely low values after rescaling, violating the diffusion equation assumptions, with unexpected impacts on population dynamics. Further analysis is needed to elucidate under what parameter combinations biases may occur in neutral forward-time simulations, or whether biases may indeed be avoidable in such simulations.

### Implications for previous simulation studies

Our findings have several implications for published population genetics studies involving forward-in-time simulations. First, simulation-based studies modelling small regions (*e.g.,* in many *D. melanogaster*-like or theoretically motivated studies) or using small scaling factors (as for many studies of human-like populations) are probably minimally biased (*e.g.,* Messer and Petrov 2013; Johri, Riall, et al. 2021). For studies simulating longer regions with moderately high scaling factors, specific summary statistics such as diversity may be reduced while other population genetic statistics such as the SFS and LD patterns are expected to be preserved (*e.g.*, Marsh and Johri 2024). Finally, studies that have simulated either a single selective sweep or recurrent sweeps in small functional regions are unlikely to be biased due to rescaling (*e.g.,* Elyashiv et al. 2016; Booker and Keightley 2018).

Simulation-based inference approaches like ABC (Beaumont et al 2022) often employ large scaling factors because ABC is computationally very intensive and simulations with large ancestral population sizes due to wide priors can be time-consuming. However, as these simulations have usually involved relatively short regions (*e.g.,* Uricchio et al. 2019; Johri et al. 2020; Johri et al. 2023), they are unlikely to be affected by rescaling. Moreover, the deviation from unscaled scenarios in many cases are so minor (e.g. Figure 2, Figure 3, Table 2) that it is unclear if it would impact inference using ABC-like methods that are dealing with bigger issues like the curse of dimensionality. In other words, inference using ABC involves optimizing the fit to multiple summary statistics and thus minor deviations are unlikely to drastically bias inference.

### General recommendations for tractable forward-simulations

Generally, we recommend designing rescaled simulations with scaling factor and length combination values that keep *R* low enough to avoid high rates of multiple crossovers, unless there is evidence of common multiple crossovers in the target species. The probability of multiple crossovers occurring in each individual per generation, *P*(> 1 *CO*), is only dependent on *R* when assuming no crossover interference (as in *SLiM*). *P*(> 1 *CO*) can be calculated from the probabilities of 0 crossovers per individual, *P*(0 *CO*), and 1 crossover per individual per generation, *P*(1 *CO*), as 1 − *P*(0 *CO*) − *P*(1 *CO*) which equates to:

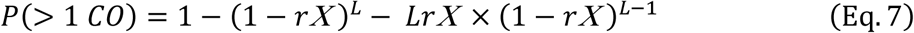

An *R* value of 0.44 corresponds to *P*(> 1 *CO*) of approximately 5% and may be a useful upper limit when trying to avoid high rates of multiple crossovers and their related effects. Note that when multiple crossovers are common in the modelled population, it may be necessary to model multiple crossovers using forward-in-time simulators or consider alternative implementations to better model realistic recombination dynamics with interference. Finally, users should account for frequent multiple crossovers using appropriate mapping functions (*e.g.,* Haldane 1919) with rescaled parameters when comparing to analytical predictions, and consider the effects that a non-linear map may have on their results.

The values of *U* and *R* needed to avoid pervasive HRI and progeny skew, when present at a much lower extent in the unscaled population, will depend heavily on the distribution of fitness effects and the proportion of selected sites which are beneficial. HRI is difficult to assess directly, and often it may be most pragmatic to simply test a range of *X* × *L* values. Typically, the optimal values for a given analysis should ensure that minor deviations in *U* and *R* do not change the observed diversity, SFS, LD patterns, or fixation probabilities of selected mutations calculated from simulated data, unless expected in the real population. It is possible to derive more direct statistics for interference between beneficial mutations such as the probability of a beneficial mutation arising while another segregates in the population (see for example *P*_*inter*_ in Uricchio and Hernandez 2014). Daigle and Johri (2024) have suggested that values of 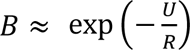 can themselves be very informative about the extent of HRI in the simulated population and that values below 0.25 might lead to severe effects. Thus, if the *B* values obtained from simulations are a lot lower than those predicted theoretically for the unscaled population, it might be worth decreasing the scaling factor (as shown in Table 1). Progeny skew should be monitored in rescaled populations by checking the distribution of offspring counts per parental haplotype to ensure it is either Poisson-distributed and follows Wright-Fisher assumptions, or so that it matches the expected distribution of the modelled population.

Finally, many replicates may be necessary to provide sufficient information to overcome stochasticity when simulating short regions, and directional biases may occur when modelling very short regions (≤ 10 kb). As a result, for *D. melanogaster* simulations under most selection parameters we would recommend regions of length *L* = 200 kb (equivalent to ∼28 genes), with a scaling factor *X* < 250 and at least 10 replicates as appropriate for many simulation-based population genetics analyses to avoid the unintended biases due to rescaling that we observed.

### Conclusion

Our results expose the limits of effective rescaling when simulating long genomic regions, which may restrict current research seeking to realistically model genome-wide effects of selection. Moreover, as unlinked effects of background selection do not scale with *N* (Santiago and Caballero 1998; also see Appendix of Charlesworth 2012), it’s unclear how one can simulate a full genome with multiple chromosomes experiencing selection simultaneously with rescaling (Matheson and Masel 2024). Thus, it may be best to limit forward simulations with selection to individual chromosomal regions (if rescaling is involved) and perhaps attempt to account for unlinked effects of background selection by a simple rescaling of *N*_*e*_; such considerations, however, need further testing.

Despite our focus on pitfalls and limitations associated with highly rescaled forward-in-time simulations, it is important to highlight our null results, which demonstrate the effectiveness of rescaling in modelling sequence evolution of complex populations in a tractable manner, given appropriate informed parameter choices. Furthermore, with rapid recent developments in simulation software and programming interfaces there is great potential for advancements in how forward-in-time simulators model complex population genetics dynamics associated with recombination and selection. Finally, we hope that our findings contribute to the ongoing development of robust frameworks and standards to help facilitate effective, unbiased population genetics analysis with forward-in-time simulations.

## Acknowledgements

The research in this study was conducted using computational resources provided by ITS Research Computing at the University of North Carolina at Chapel Hill. The authors declare no conflicts of interest. We thank Brian Charlesworth, Jeff Jensen and Fanny Pouyet for feedback on drafts that greatly improved the manuscript. We also thank Brian Charlesworth and Jeff Jensen for insightful comments regarding the effects of multiple crossovers and progeny skew respectively. Research reported in this publication was supported by the National Institute of General Medical Sciences of the National Institutes of Health under award number R35GM154969 to PJ.

## Data availability

All relevant code and scripts generated for this study are available at https://github.com/JohriLab/Simulation_Rescaling.

